# Minimally-invasive Manipulation of Spared and Hypoactive Interneurons reduces CA1 Synchronization and Nonspatial Behavior alterations in Epilepsy models

**DOI:** 10.1101/2024.08.02.606307

**Authors:** Célanie Matringhen, Alexandre Vigier, Nikoleta Bourtouli, François J. Michel, Thomas Marissal

## Abstract

Temporal lobe epilepsy (TLE) is associated with seizures and severe cognitive impairments including memory deficits. The dysfunction of a large population of hippocampal interneurons, composed of various subtypes, is proposed as a key mechanism and a therapeutical target. However, the nature and extent of alterations in hippocampal inhibitory neurons remain unclear, as does their impact on TLE pathology. To address this issue, we combined immunolabeling, calcium imaging, electrophysiology, chemogenetic tools, and behavioral assays in mouse pilocarpine TLE models to investigate the survival and changes in the activity of a large population of Dlx-expressing interneurons and test the effect of their manipulation on seizures and cognitive impairments. We observed that a large population of Dlx-expressing interneurons, that includes Cholecystokinine-, Parvalbumin- and Somatostatin-positive cells, were overall spared from histological damage occurring in CA1 from TLE mice, despite subtype-specific vulnerability. In addition, Dlx-expressing interneurons exhibited reduced activity *in vitro*, correlated to hypersynchrony in the CA1 network. Enhancing CA1 interneuron discharge *in vitro* using a chemogenetic strategy rescued CA1 activity and synchronization. *In vivo*, a minimally invasive strategy to normalize interneuron activity does not rescue the full range of pathological features associated with TLE, but significantly reduces some cognitive impairments, such as behaviors related to nonspatial learning and memory. Our findings suggest that enhancing local CA1 interneuron activity can restore local network balance and improve cognitive function in TLE.

## 1. Introduction

Temporal lobe epilepsy (TLE) is a prevalent and often incurable disease associated with the generation of spontaneous recurrent seizures as well as severe cognitive impairments including memory deficits (Marissal, 2021). The dysfunction of hippocampal interneurons is proposed as a key mechanism and a therapeutical target (Liu et al., 2014). However, the nature and extent of alterations in hippocampal interneurons remain controversial despite an abundant literature (Cossart et al., 2001; Dinocourt et al., 2003; Peng et al., 2013; Proddutur et al., 2023; Shuman et al., 2020), as does the validity of interneuron-specific strategies designed to reduce the seizures and cognitive impairments related to TLE. Various strategies based on stem cell grafting or reactive glia reprogramming have been tested to replenish the pool of hippocampal interneurons (Bershteyn et al., 2023; Lentini et al., 2021), which are postulated to undergo a subclass-specific loss during the early disease stages according to pioneer studies in the CA1 area from TLE rats (Dinocourt et al., 2003). However, the clinical application of these approaches remains questionable given that recent observations from patients and mouse models of TLE suggest that the loss of CA1 inhibitory neurons can be modest or absent, and unlikely to contribute to behavioral coding deficits (Lehner et al., 2024; Shuman et al., 2020). Other approaches are based on the specific manipulation of a single class of interneurons (e.g., Parvalbumin or PV neurons), using chemogenetic or optogenetic tools, sometimes combined with a closed-loop system with EEG monitoring. Again, apart from the difficulties of implementing such strategies in patients, these approaches also face several limitations. Firstly, most are based on the excitation of an interneuron subtype postulated to be already hyperactive or hyperexcitable in epileptic conditions (Cossart et al., 2001), and their effect on seizures (Krook-Magnuson et al., 2013; Lévesque et al., 2019; Wang et al., 2020) or behavioral alterations (Kim et al., 2020; Wang et al., 2018) is not always beneficial. Secondly, these studies overlook the extent of degeneration of the considered interneuron subtype, a factor that could nonetheless affect the effect of their manipulation. Lastly, these studies rarely take into account the complementary pathophysiological role played by the different classes of interneurons expressing PV, Somatostatin (SOM) or Cholecystokinin (CCK) (Amilhon et al., 2015; Dudok et al., 2022, 2021; Miri et al., 2018; Parrish et al., 2019). These different interneuron subclasses form interconnected inhibitory microcircuits that regulate CA1 network dynamics (Bocchio et al., 2024), and activation of a large population of interneuron could control pathological activity more efficiently than stimulation of a single subpopulation (Ledri et al., 2014). To tackle these issues, we combined immunolabelling and in vitro calcium imaging in mouse TLE models to investigate the survival and activity changes of a large population of Dlx-positive CA1 interneurons, including inhibitory neurons expressing PV, SOM and CCK. Then, we tested the effect of manipulating this population of Dlx-expressing CA1 interneurons on TLE-related seizures and cognitive alterations using behavioral assays, *in vivo* electrophysiology and minimally invasive chemogenetic tools that are potentially applicable to humans (Dimidschstein et al., 2016). We show that a large population of Dlx-expressing interneurons, that includes CCK-, PV- and SOM-positive cells, were overall spared from CA1 atrophy observed in the TLE mice, despite differential subtype resistance. Overall, CA1 Dlx-expressing interneurons are less active in epileptic than in control conditions *in vitro*, resulting in increased activity and synchronization of the epileptic CA1 network. Enhancing CA1 Dlx-expressing interneuron discharge using a chemogenetic strategy *in vitro* rescued CA1 activity and synchronization. *In vivo*, the minimally invasive chemogenetic activation of hippocampal Dlx-expressing interneurons does not affect seizures nor deficits related to spatial learning and memory, but does reduce impairments in behaviors related to nonspatial learning and memory. Therefore, we propose that rescuing CA1 local network dynamics using interneurons as a lever could be sufficient to decrease specific behavioral deficits.

## 2. Materials and Methods

To study whether CA1 interneurons are major actors and potential therapeutic target in TLE pathology and particularly behavioral impairments, we used a combination of immunolabeling, two-photon calcium imaging, electrophysiological recordings, chemogenetic tools, and behavioral testing performed in mouse pilocarpine-based models *in vitro* (Goirand-Lopez et al., 2023) and *in vivo* (Vigier et al., 2021). All experiments conducted on mice were performed in accordance with the directives of European Community Council (2010/63/UE) and received approval from the French Ministry for Research, after ethical evaluation by the institutional animal care and use committee of Aix-Marseille University (APAFIS number: #23627).

The procedures are detailed in the Supplemental Information: <u>*Animals,* in vivo *and* in vitro *pilocarpine-based models of TLE, Surgeries*</u>, <u>*Behavioral testing*</u>, <u>*Histology*</u>, <u>*Calcium imaging and analysis*</u>, <u>*EEG recordings and analysis*</u>, and <u>*Statistical methods*</u>.

## 3. Results

### 3.1. Dlx-expressing interneurons are overall spared in the CA1 from epileptic mice

To study the fate of a large population of CA1 interneurons in epileptic mice, we injected non-epileptic control mice and pilocarpine-treated epileptic animals (“Pilo”) with AAVs expressing the fluorescent reporter tdTomato into cells expressing the interneuron-specific enhancer Dlx (see **Material and Methods**) (Dimidschstein et al., 2016). The population of Dlx-expressing interneurons, including PV-, SOM- and CCK-positive inhibitory cells (**Fig. 1A**), showed no significant differences in density or distribution across the different CA1 sublayers in the epileptic mice compared with controls (**Fig. 1B-D**), despite histological alterations similar to those observed in patients (i.e., CA1 atrophy associated with hippocampal sclerosis, **Supplemental Fig. 1A-B**) (Costa et al., 2019). As expected, the overall survival of Dlx interneurons masked differential vulnerability depending on the major inhibitory cell subtype (**Supplemental Fig. 1C-G**). In line with (Shuman et al., 2020), our data did not support a loss of CA1 PV and SOM interneurons in epileptic mice (**Supplemental Fig. 1C, E-F**). In contrast, CCK interneurons were significantly decreased in pilocarpine-treated mice compared with controls (**Supplemental Fig. 1 D, G**). The distribution of PV, SOM and CCK interneurons remained unchanged (**Supplemental Fig. 1 E-G**).

**Fig. 1:**
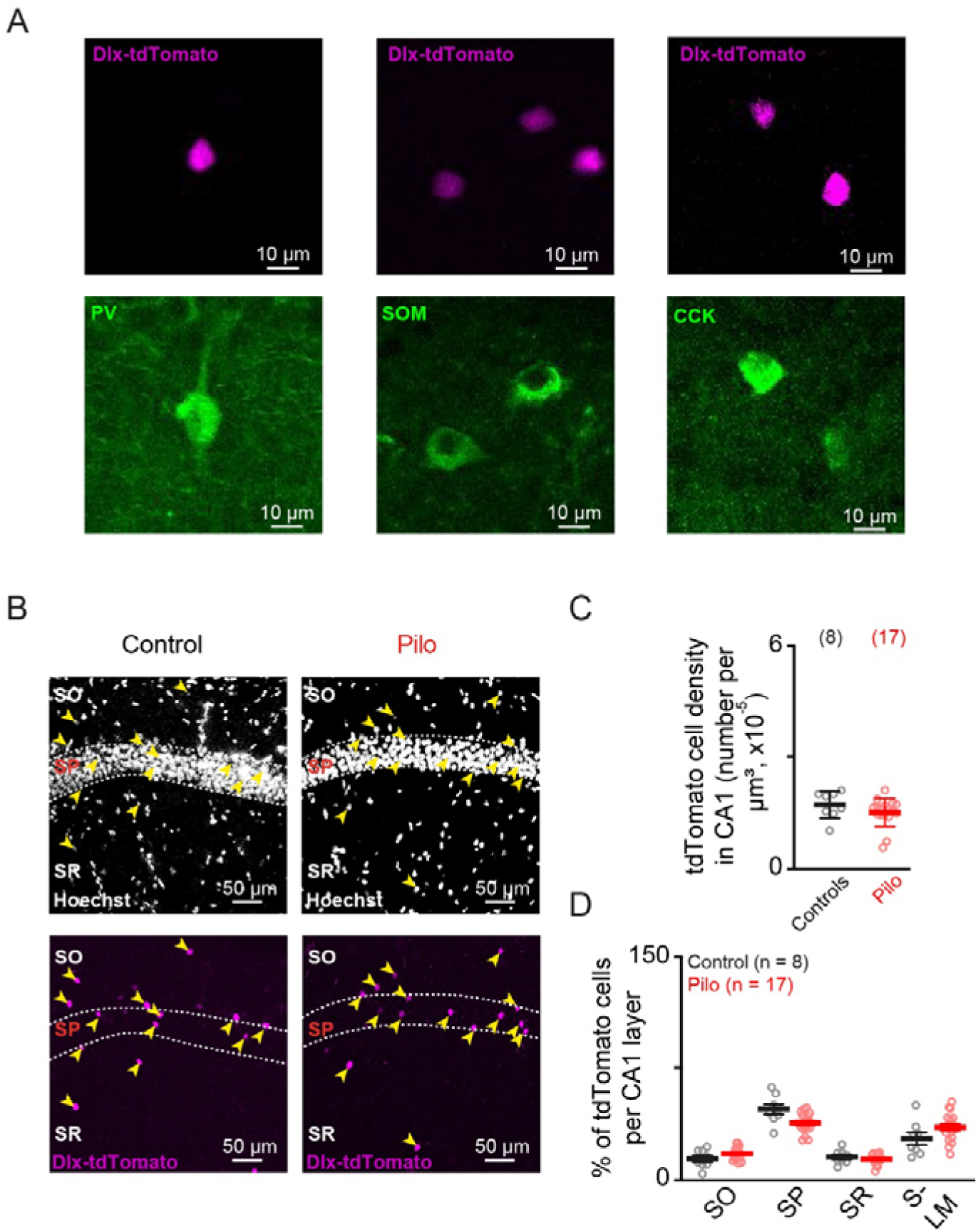
Dlx-expressing interneurons are overall spared in TLE mice. **A.** Examples of CA1 Dlx-expressing interneurons that co-express tdTomato (top) and PV (bottom left), SOM (bottom middle), or CCK (bottom right) in the hippocampus infected with AAV9-hDlx-GqDREADD-tdTomato-Fishell-4. **B.** Representative Hoechst staining (top), as well as tdTomato labelling (bottom) in the CA1 region from a control animal (left) or a pilocarpine-treated epileptic mouse (Pilo, right) infected with AAV9-hDlx-GqDREADD-tdTomato-Fishell-4 in the hippocampus. Neurons with Hoechst and tdTomato co-labelling are indicated by a yellow arrow. **C.** Number of tdTomato-labelled interneurons per µm^3^ of CA1. Two-tailed Mann-Whitney test *P =* 0.2107. **D.** Percentage of tdTomato-labelled interneurons per CA1 layer (two-way ANOVA experimental condition effect F (2, 36) = 0.8553, *P =* 0.4336). SO: Stratum Oriens, SP: Stratum Pyramidale, SR: Stratum Radiatum, SLM: Stratum Lacunosum-Moleculare. Data are presented as mean ± s.e.m. Each circle represents a given mouse (The number of mice is specified in parentheses).

In conclusion, our results suggest that a substantial number of Dlx-expressing interneurons survive the epileptic insult despite histological alterations and subtype-specific degeneration in the CA1 from Pilo mice.

### 3.2. Hypersynchronization of CA1 network in vitro is correlated with hypoactive interneurons and is reversible

Numerous studies based on different models suggest that the different categories of CA1 inhibitory neurons surviving TLE could display altered activity. However, the directionality of these alteration (i.e., hyper- or an hypo-activity), and whether their activation could produce a positive or negative effect on the pathological traits associated with TLE remains unclear (Dudok et al., 2022). To tackle this issue, we monitored the calcium activity of a large number of CA1 neurons as in (Marissal et al., 2018), using *in vitro* GCaMP8m-based calcium imaging in control versus pilocarpine-treated slices (Controls versus Pilo, **Fig. 2A-E**), which exhibit physiological and histological alterations similar to those observed in the hippocampus of pilocarpine-treated mice or even in hippocampal tissues from adult TLE patients (Boileau et al., 2023; Peret et al., 2014). The calcium activity of CA1 was recorded under the continuous presence of 50µM carbachol (Cch), which induces in control conditions network activity resembling those that occur *in vivo* during navigation, learning and memory processes (Fisahn et al., 1998). As expected (Li et al., 2019; Marissal et al., 2018), the Cch-induced CA1 calcium activity reflects the neuronal action potential discharge, given that most calcium transients were abolished in the presence of the sodium channel blocker TTX (**Supplemental Fig. 2A-C**).

**Fig. 2:**
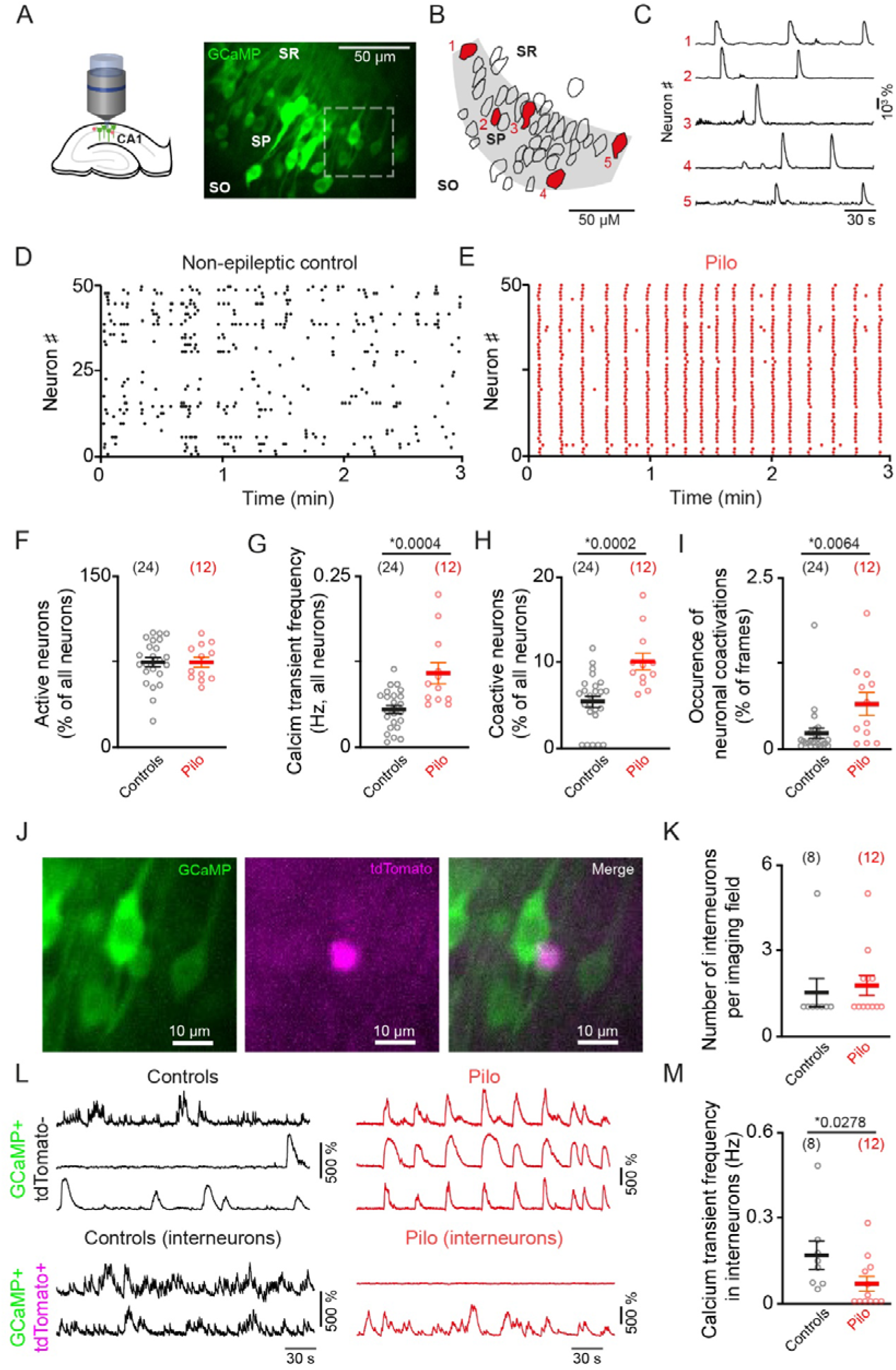
Epileptic CA1 network is hyperactive and synchronous while Dlx interneurons are less active *in vitro*. **A.** Schematic representation of the experimental procedure (left) and photograph of GCaMP8m-expressing neurons in the hippocampal CA1 region (right). SO: Stratum Oriens, SP: Stratum Pyramidale, SR: Stratum Radiatum. **B.** Contours of GCaMP8m-expressing neurons shown in A. **C.** Calcium transients recorded in the presence of carbachol (50 μM) from selected neurons identified as 1 to 5 in B (represented with red-filled contours). **D-E.** Representative raster plots of calcium transients recorded in control (D, black) or Pilo (E, red) conditions. **F.** Proportions of CA1 neurons displaying at least one calcium transient during the duration of the movie in each condition (Two-tailed unpaired t-test, *P =* 0,9844). **G.** Frequency of calcium transients recorded in all CA1 neurons (Two-tailed unpaired t-test). **H-I.** Percentage of co-active neurons above chance level (H) and occurrence (I) of these co-activations (H, I: Two-tailed Mann-Whitney test). **J.** Photograph of the field represented by a dashed rectangle in A, in which a Dlx-expressing interneuron co-express GCaMP8m and tdTomato. **K.** Number of interneurons per calcium imaging field (Two-tailed Mann-Whitney test, *P =* 0,3246). **L.** Examples of calcium activity exhibited by GCaMP+ expressing tdTomato-negative CA1 neurons (top) or tdTomato-positive interneurons (bottom) in control (left) or pilo (right) conditions. Each trace represents a different neuron. **M.** Average frequency of calcium transients recorded in CA1 interneurons per slice (Two-tailed Mann-Whitney test). Asterisks indicate significant differences (*P <* 0.05). Data are presented as mean ± s.e.m. Each circle represents the mean for a given slice (the number of slices is specified in parentheses. Slices were prepared from *N =* 7 and 5 animals in control and Pilo conditions, respectively).

While the percentage of active cells was similar between both conditions (**Fig. 2F**), CA1 neuron calcium activity was more frequent and synchronous in Pilo conditions than in control slices, which was reflected by significantly higher values in the frequency of calcium transients (**Fig. 2G**), the number of coactive neurons (**Fig. 2H**) as well as the occurrence of these neuronal coactivations (**Fig. 2I**). We reported no difference between experimental groups in the average amplitude and duration of calcium transients (**Supplemental Fig. 2D-E**).

We also specifically recorded the GCaMP8m-based calcium activity of Dlx-expressing interneurons (see **Materials and Methods**, **Fig. 2J-M**). Overall, Dlx-expressing CA1 interneurons seemed to be spared from the histological alterations that affect pilocarpine-treated slices (Peret et al., 2014), given that the number of interneurons per calcium imaging field was not different between conditions (**Fig. 2K**), while the number of GCaMP-positive neurons per field was decreased in Pilo condition (**Supplemental Fig. 2F**). By contrast, the calcium transients generated by CA1 interneurons were significantly less frequent under Pilo conditions, or even abolished completely (**Fig. 2L, M** and **Supplemental Fig. 2G**). Importantly, interneuron calcium activity was not significantly affected by their localization in deep CA1 layers (i.e., strata oriens and pyramidale) where most PV and SOM neuron soma are situated, or superficial CA1 layers (i.e., strata radiatum and lacunosum-moleculare), where most CCK cell bodies are localized (**Supplemental Fig. 2G).**

In conclusion, our data suggest that while the CA1 neuronal network is more active and synchronous, interneurons are significantly less active in Pilo conditions compared to control slices.

Could an interneuron-targeting chemogenetic strategy restore the activity of CA1 interneurons and re-establish control-like network dynamics? To test this, we compared the *in vitro* calcium activity in the Pilo CA1 neuronal network, including a few tdTomato- and hM3D-expressing interneurons, before (‘Pilo-VEH’) and after 15 minutes of bath application of CNO (Clozapine-N-Oxide, 2 µM, see **Materials and Methods)**. The activation of the excitatory DREADD hM3D using CNO significantly increased the frequency of calcium transient generated by inhibitory neurons (**Fig. 3A-B**), to levels that are comparable to the Controls (Mann-Whitney test *P =* 0.5148, **Fig. 2J-M**).

**Fig. 3:**
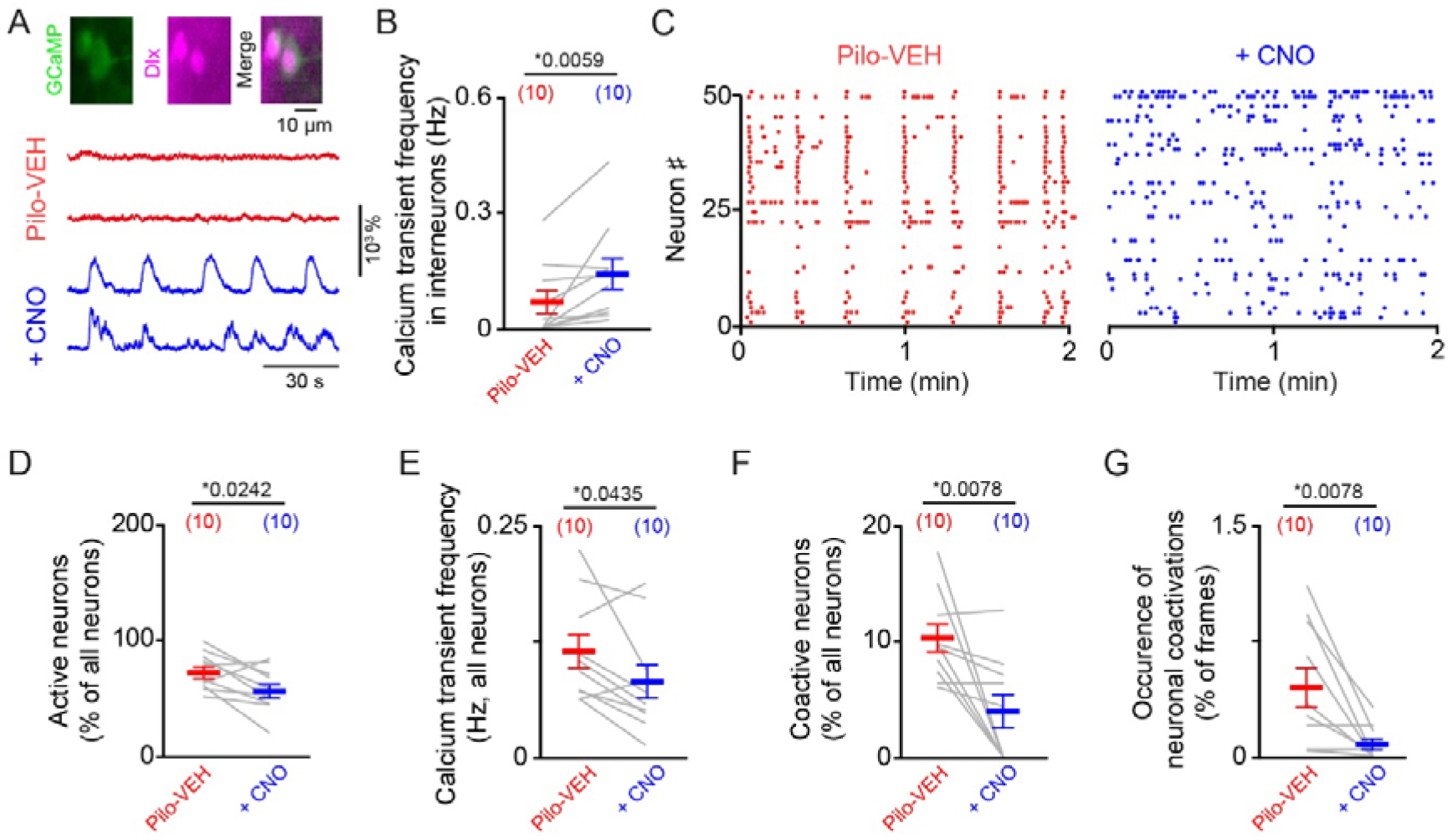
Reestablishing Dlx interneuron activity decreases local CA1 network synchronization *in vitro*. **A.** Photographs of CA1 interneurons that co-express GCaMP8m and tdTomato (fused to hM3D) (top) and example of calcium activity exhibited by hM3D-expressing CA1 interneurons in Pilo condition before (red, Pilo-VEH) and after 15 minutes of treatment with CNO (2 µM, blue) (bottom). **B.** Frequency of calcium transients recorded in hM3D-expressing CA1 interneurons in Pilo condition before and after treatment with CNO (Two-tailed Wilcoxon test). **C.** Representative raster plots of calcium transients recorded in the same slice before and after treatment with CNO. **D-G.** Levels of calcium activity (D, E) and synchronization (F, G), reflected by the number of active neurons (D), the frequency of calcium transients recorded in all CA1 neurons (E), the percentage of co-active neurons above chance level (F) and the occurrence of these co-activations (G) in both conditions (Two-tailed paired t-test for D and E, Two-tailed Wilcoxon test for F and G). Data are presented as mean ± s.e.m. Each gray line represents a given slice (the number of slices is specified in parentheses. Slices were prepared from *N =* 3 animals).

The chemogenic elevation of CA1 interneuron discharge attenuated the Pilo CA1 network activity and synchronization (**Fig. 3C**), which was reflected by a significant decrease in the percentage of active cells (**Fig. 3D**), the average frequency of calcium transients generated by all neurons of the imaged fields (**Fig. 3E**), the number of neurons involved in coactivations (**Fig. 3F**), and the occurrence of these coactivations (**Fig. 3G**). By contrast, the chemoactivation of hippocampal interneurons affected neither the amplitude nor the duration of calcium transients (**Supplemental Fig. 2H, I**). Apart from the number of active cells, the *in vitro* calcium activity in pilocarpine-treated CA1 treated with CNO was no longer significantly different from the control slices (Two-tailed unpaired t-test *P =* 0.0331, 0.0820, 0.4058, and 0.2157 for the number of active cells, as well as calcium transient frequency, amplitude and duration, respectively; Mann-Whitney test *P =* 0.4669 and 0.1678 for the for the number of coactive neurons and the occurrence of coactivations, respectively. **Figure 2F-I**). Therefore, our data suggest that rescuing interneuron activity is sufficient to restore local CA1 network dynamics *in vitro*.

### 3.3. The in vivo chemoactivation of hippocampal interneurons does not rescue the pathology, but attenuates specific cognitive deficits

Could the chemogenetic stimulation of hippocampal Dlx-expressing interneuron activity decrease the pathological traits associated with TLE, such as seizures and cognitive deficits?

To answer this question, we first performed *in vivo* EEG recordings on pilocarpine-treated TLE animals while hM3D receptors, specifically expressed in Dlx-positive interneurons, were selectively activated through the presence of CNO in the drinking water (‘Pilo-CNO’ condition, estimated daily consummation for EEG experiments: 7.6 mg/kg of CNO per mouse and day), or not (Pilo-VEH condition, see **Materials and Methods**). Importantly, the density and distribution of Dlx-positive interneurons (that express hM3D receptors and tdTomato) was similar for all experimental conditions (**Supplemental Fig. 3A-C**). As expected, prolonged ictal activity reminiscent of generalized tonic-clonic seizures has been monitored in all pilocarpine-treated animals in the hippocampal region distant from injection sites (**Fig. 4A**, see **Materials and Methods**). Despite the impact of interneuron chemoactivation on local synchronization *in vitro* (**Figure 3**), we observed no significant difference on the frequency (**Fig. 4B**) and duration (**Supplemental Fig. 3D**) of seizures between Pilo-VEH and Pilo-CNO conditions.

**Fig. 4:**
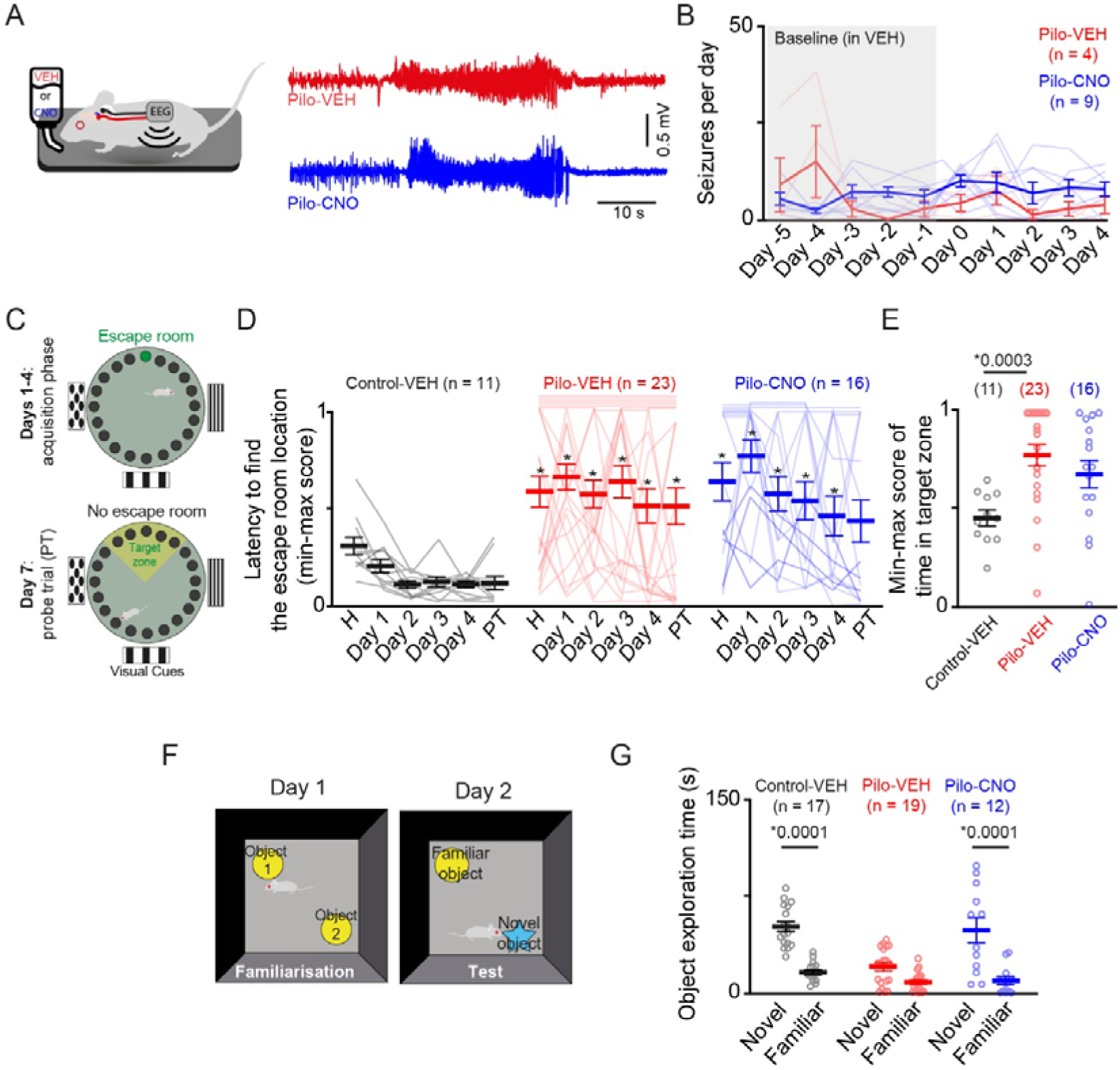
The *in vivo* chemoactivation of Dlx interneurons do not rescue TLE pathology, but reduces nonspatial behavioral deficits. **A.** Schematic representation of the experimental procedure (top) and example traces of ictal activity recorded with telemetric EEG recording in freely moving epileptic mice in the absence (Pilo-VEH) or presence of CNO (Pilo-CNO) in the drinking water to specifically elevate or not the activity of Dlx- and hM3D-expressing hippocampal interneurons (bottom). **B.** Frequency of ictal events in Pilo-VEH versus Pilo-CNO conditions across days of recording (Two-way ANOVA experimental group effect F (1, 11) = 1.338 *P =* 0.2719. **C.** Schematic representation of the Barnes Circular Maze (BCM) acquisition phase (top) and probe trial (bottom). **D.** Min-max scores of latencies to find the escape room during the BCM acquisition phase. Data were compared in Control-VEH versus Pilo-VEH groups (Two-way ANOVA experimental group effect F (1, 32) = 23.10 *P <* 0.0001, followed by Sidak’s post hoc test, *P =* 0.0251 0.0001, 0.0001, 0.0001, 0.0009 and 0.0030 for habituation, acquisition days 1-4 and PT, respectively), Control-VEH versus Pilo-CNO groups (Two-way ANOVA experimental group and treatment effect F (1, 25) = 29.67 *P <* 0.0001, followed by Sidak’s post hoc test, *P =* 0.0371 0.0001, 0.0005, 0.0040, 0.0221 and 0.0715 for habituation, acquisition days 1-4 and PT), and Pilo-VEH versus Pilo-CNO groups (Two-way ANOVA treatment effect F (1, 37) = 0.0161, *P =* 0.8999). Straight horizontal lines across the top of the plots indicate that a mouse has scored 1 for two or more consecutive days. **E.** Min-max scores of times spent in the target zone during the BCM probe trial in various experimental conditions (Two-tailed Mann-Whitney test for the comparison between Control-VEH and Pilo-VEH and between Pilo-VEH and Pilo-CNO *P =* 0.1789, unpaired t-test for the comparison between control-VEH and Pilo-CNO, *P =* 0.0230). **F.** Schematic representation of the NORT procedure. **G.** Time spent exploring the novel and the familiar object during the NORT test phase, in each experimental condition (two-way ANOVA interaction experimental condition x type of object F (2, 45) = 8.182, *P =* 0.0009, followed by Sidak’s post hoc test to compare the time in novel versus familiar object, *P <* 0.0001 in Control-VEH and Pilo-CNO conditions, *P =* 0.0508 in Pilo-VEH condition). Asterisks indicate significant differences. Data are presented as mean ± s.e.m, unless specified otherwise. Each circle or line represents a given mouse (The number of mice is specified in parentheses).

In parallel, we performed behavioral testing on pilocarpine-treated TLE animals with or without chemoactivation of Dlx-expressing hippocampal interneurons. We first used the Barnes Circular maze (BCM) to assess behavior related to spatial learning and memory in epileptic mice while hM3D-expressing hippocampal interneurons were selectively activated through the presence of CNO in the drinking water (i.e., ‘Pilo-CNO’ condition), or not (‘Pilo-VEH’ condition). We also used a group of vehicle-treated non-epileptic control mice (‘Control-VEHs’).

Animals in the Pilo-VEH group show impaired behavior in the BCM compared to Control-VEH mice (**Fig. 4C-E**, **Supplemental Fig. 3E-G, Supplemental Fig. 4A**). Notably, the min-max scores related to the latency to locate the escape room at any BCM stage (acquisition days 1-4 and probe trial or PT, **Fig. 4D**), and the time in the target zone during the PT (**Fig. 4E**) were significantly higher in Pilo-VEH mice, reflecting poorer performance (**Material and Methods**). The proportion of Pilo-VEH animals that identified the location of the escape room during all days of BCM (success rate, **Supplemental Fig. 3E-F**) and exhibited direct exploration strategy (**Supplemental Fig. 3G**) is significantly lower than in Control-VEH animals. Last, Pilo-VEH mice display highest min-max scores related to number of primary errors (**Supplemental Fig. 3G**). The original data is indicated in **Supplemental Fig. 4A**. Epileptic mice treated with CNO (estimated consumption for behavioral experiments : 6.4 mg/kg per mouse and day) display little enhancement of their performance in the BCM (**Fig. 4C-E**, **Supplemental Fig. 3E-G, Supplemental Fig. 4A**) despite a slight improvement of their min-max scores related to the latency to locate the escape room (**Fig. 4D**) and success rates (**Supplemental Fig. 3E-F**) during the later phases of the test (e.g., acquisition day 4 and PT).

However, the analysis of BCM was complicated by the multitude of aberrant behaviors exhibited by the epileptic mice (e.g., jumping off the apparatus, no movement from the release zone) (see **Materials and Methods, Supplemental Fig. 4A**), which were not necessarily related to spatial processes, or even to CA1 circuits. To investigate other types of CA1-dependent memory that are altered in TLE, we conducted novel object recognition task (NORT) on all experimental groups (see **Materials and Methods, Fig. 4F-G**). During the NORT familiarization phase, no experimental group showed a preference for either of the two identical objects (**Supplemental Fig. 4B**). During the test phase, the Control-VEH mice explored the novel object significantly more than the familiar object while Pilo-VEH mice spent a similar amount of time exploring the two objects in the absence of Dlx-expressing interneuron manipulation. By contrast, preference for the novel object was rescued in Pilo-CNO animals (**Fig. 4G**). Regarding locomotory activity, although Pilo animals displayed a greater variability compared to Controls, we observed no significant group differences in the distance covered during the BCM probe trial, or the NORT familiarization and test phases (**Supplemental Fig. 4C**).

Our data suggest that chemoactivation of a large population of Dlx-expressing hippocampal interneurons does not rescue the full range of pathological features associated with TLE, since seizures and deficits related to spatial learning and memory are unaffected. However, activation of Dlx-expressing interneurons in the hippocampus does reduce some cognitive impairments, such as behaviors related to nonspatial learning and memory.

## 4. Discussion

Using immunohistological analysis, we show that pilocarpine-treated TLE mice display no decrease in the density of CA1 Dlx-expressing interneurons, a large population of inhibitory cells that includes PV-, SOM- and CCK-expressing interneurons. The overall resistance of Dlx-expressing interneurons hides diversity in the vulnerability of inhibitory neuron subtypes. Surprisingly, we showed an increase in CA1 PV- and SOM-expressing interneuron density in pilocarpine-treated TLE mice. Our observations suggest that this phenomenon is more likely explained by the decrease in CA1 area (reflecting CA1 atrophy associated with hippocampal sclerosis)(Costa et al., 2019) than by a plasticity mechanism (Donato et al., 2013). Although in contradiction with some pioneer observations in the epileptic rat (Cossart et al., 2001; Dinocourt et al., 2003), these data are in agreement with recent studies performed on tissues from patients, as well as others performed using the mouse pilocarpine model, which suggests that spatial processing impairment is not correlated to a loss of CA1 PV and SOM interneuron (Lehner et al., 2024; Shuman et al., 2020). In contrast, CCK interneurons are diminished in epileptic mice, as in other models of TLE (Kang et al., 2021). Given that the silencing of CA1 CCK interneurons does not impact mouse performances in a spatial memory test comparable to the BCM (Guerrero et al., 2024), we do not believe that the reduction of their density alone can explain the cognitive alterations displayed by pilocarpine-treated mice.

In line, our *in vitro* calcium imaging data suggest that Dlx-positive interneurons are not diminished in pilocarpine-treated slices compared to controls, despite a smaller number of GCaMP-expressing neurons, which could be the consequence of *in vitro* histological alterations similar to those observed *in vivo* in TLE mice or even in patients (Boileau et al., 2023; Peret et al., 2014). Dlx-expressing interneurons are overall hypoactive under epileptic conditions *in vitro*, while the network presents excessive and hypersynchronous activity. This observation brings new insight to the debate as to whether hippocampal interneurons are less active (“dormant”) or rather hyperactive. We did not attempt to decipher the mechanisms that explain the reduced activity of CA1 interneurons *in vitro*. Several studies suggest that the intrinsic and synaptic properties of certain inhibitory neurons may be altered under epileptic conditions, thereby permanently modifying their firing (Dugladze et al., 2007; Lee et al., 2024; Proddutur et al., 2023). However, characterizing the pathophysiological properties of Dlx-expressing interneuron population, which includes various morpho-physiological subtypes (Dimidschstein et al., 2016), would be an entire study on its own. Similarly, future investigations are needed to determine whether the activity of hippocampal interneuron subtypes is also altered *in vivo*, while epileptic mice are behaving. Interestingly, our data shown in **Supplemental Fig. 2G** suggest that the reduction of activity is more marked for interneurons localized in the superficial CA1 layers (i.e., strata radiatum and lacunosum-moleculare), which includes CCK-positive cells. In addition, our result suggest that, despite the average decrease of interneuron activity, a small subpopulation of inhibitory cells localized close to the pyramidal layer is hyperactive as in (Cossart et al., 2001).

As this study focused on the activity of Dlx-expressing interneurons in controls or under conditions reproducing the alterations associated with TLE, we did not combine *in vitro* calcium imaging with local field potential recordings to characterize the activity patterns generated by hippocampi treated or not with pilocarpine. Under control conditions, carbachol-induced GCaMP fluorescence variations in CA1 correlate with physiological activities such as theta rhythms (Li et al., 2019). In the pilocarpine-treated hippocampus, which shows similar alterations to those observed *in vivo* or in patients, excessive and hypersynchronous calcium activity could be explained by the abnormal recruitment of CA1 neurons into altered rhythmic activities or interictal-like-epileptiform events (Goirand-Lopez et al., 2023). Either way, chemogenetic activation of Dlx-expressing interneurons restores CA1 network dynamics towards control levels.

Most, if not all, studies testing the effect of manipulating a subclass of interneurons on the pathological features of TLE overlook the nature and extent of alterations affecting the neuronal type of interest. Therefore, strategies based on chemo- or optogenetic excitation of a subclass of interneurons, a substantial proportion of which may degenerate (Dinocourt et al., 2003), and the survivors of which may be hyperactive (Cossart et al., 2001; Dugladze et al., 2007), remain questionable. Furthermore, these strategies are not always beneficial for seizures (Krook-Magnuson et al., 2013; Lévesque et al., 2019; Wang et al., 2020) or behavioral alterations (Kim et al., 2020; Wang et al., 2018). In our study, we build on our prior observation that a large population of Dlx-expressing hippocampal interneurons are overall spared from histological alterations *in vivo* and display a decreased activity *in vitro*. On these premises, we used a chemogenetic strategy to enhance their discharge specifically. *In vitro*, this interneuron-specific manipulation restores the activity and synchronization of the CA1 neuronal network to levels similar to levels comparable to control. *In vivo*, the chemoactivation of Dlx-expressing interneurons does not reduce seizures recorded at a distance from the virus injection sites or spatial behavior deficits but does alleviate specific cognitive impairments such as recognizing a new object. This suggests that hippocampal interneuron manipulation does not prevent the emergence of ictal activities, which involve a broad network of brain structures (Toyoda et al., 2013). The persistence of seizures could interfere with cognition, which is only partially rescued in our conditions. However, a local normalization of hippocampal activity *in vivo*, similar to that we observed *in vitro,* could be sufficient to improve some behavior. Such normalization could be reflected by a reduction in *in vivo* epileptiform activities (e.g., interictal discharges) or the restoration of *in vivo* neuronal activities correlated with memory processes (e.g., *theta* rhythms, shown to be disrupted in pilocarpine-treated animals (Chauvière et al., 2009)). This should be the subject of further research.

We focused on the CA1 region of the hippocampus, which is considered an important node in TLE. Highly susceptible to histological damages, CA1 could not only be involved in seizure generation and propagation (Ang et al., 2006; Buckmaster et al., 2022; Wozny et al., 2005), but also in the cognitive alterations associated with TLE (Chauvière et al., 2009; Lenck-Santini and Holmes, 2008; Shuman et al., 2020), which drastically impair the patients’ quality of life (Holmes, 2013). In addition, the inhibitory microcircuits have been particularly well described in CA1 region under physiological (Klausberger and Somogyi, 2008) and pathological conditions (Cossart et al., 2001; Dinocourt et al., 2003; Dudok et al., 2022; Dugladze et al., 2007; Muldoon et al., 2015; Peng et al., 2013). However, interneuron subclasses from other key temporal lobe regions (e.g., dentate gyrus (Proddutur et al., 2023)), are also postulated to be dysfunctional in TLE and a similar study should be performed in the future to clarify the nature and extent of their specific alterations to craft the most adequate rescue strategy.

Most, if not all, studies focusing on the role of biochemically-defined inhibitory neuron subtypes in TLE are based on transgenic lines expressing a recombinase under the control of a specific promoter (i.e., PV-Cre), but also sometimes on heavy surgery to implant optical fibers for photostimulation, combined or not with an EEG device for closed-loop seizure detection (Krook-Magnuson et al., 2013; Lévesque et al., 2019). Here, we attempted to use a controllable and minimally invasive approach that could potentially be applied to patients with TLE. We used a method based on AAV vectors and regulatory sequences to restrict DREADD expression to interneurons. This method, that does not rely on the Cre-Lox system, is effective in all non-genetically tractable vertebrate species, including humans (Dimidschstein et al., 2016). The genetic modification of the hippocampus in an intractable adult patient, which would be otherwise surgically removed as last resort, is not out of the question in terms of ethics and biosafety (Ling et al., 2023; Marissal, 2021). Given that we aimed at manipulating interneuron activity for a long, uninterrupted period and with as few adverse effects as possible, we used a drinking water-based method of DREADD ligand delivery, a technique reputed to be less invasive and stressful than intra-peritoneal injections or the use of mini-pumps (Zhan et al., 2019). Although this method did not rescue all the pathology associated with TLE, it was sufficient to restore non-spatial behavior in epileptic mice, suggesting that it may be valid under certain conditions where the activity of a neuronal subtype needs to be manipulated in a sustained manner, rather than for a specific brief period (e.g., after seizure initiation) (Kim et al., 2020).

## Data Statement

The data that supports the findings of this study are available from the corresponding author, upon reasonable request.

## CRediT Author Contribution Statement

The list of authors has been established in accordance with INSERM recommendations for the signature of scientific articles in the field of life and health sciences (https://pro.inserm.fr/rubriques/recherche-responsable/integrite-scientifique/signature-des-publications-scientifiques). The author contribution to the present project was the following: Célanie Matringhen: Conceptualization, Investigation, Formal analysis, Validation, Writing - Review & Editing; Alexandre Vigier: Investigation, Formal analysis, Methodology, Writing - Review & Editing; Nikoleta Bourtouli: Investigation, Formal analysis, Writing - Review & Editing; François J. Michel: Investigation, Formal analysis, Writing - Review & Editing; Thomas Marissal: Conceptualization, Investigation, Formal analysis, Methodology, Validation, Resources, Writing - Original Draft, Visualization, Supervision, Project administration, Funding acquisition.

## Acknowledgements

The authors have benefited from some resources and facilities hosted at the INMED as well as the teams of Valérie Crépel and Ede Rancz, where the project has been initiated and completed, respectively. The authors want to thank Ma Xiao, Emilie Pallesi, Severine Pellegrino, Philippe Moudery, Céline Boileau, Catherine Faivre-Sarrailh, Jean-Bernard Manent, Charles Quairiaux, Pierre-Pascal Lenck-Santini, Elodie Fino, Agnès Baude, Marie Kurz for having led their respective facilities, contributed to student supervision, and/or provided technical or administrative help and useful discussions. Part of the data was presented as a pre-print (DOI: 10.1101/2024.08.02.606307).

## Funding sources

This work was supported by the Institut National de la Santé et de la Recherche Médicale (INSERM) and their First Step and International Research Project programs, Aix-Marseille University (AMU), the Agence Nationale de la Recherche, the Foundation A*Midex (AMX-22-RE-AB-161), the Foundation Fyssen, the Neuroschool MSc incoming mobility program, the NeuroMarseille International Collaborative Research program (NM2MARIINM), the Fondation pour la Recherche sur le Cerveau (FRC), the PHC-Germaine-de-Staël program and the Ligue Française contre l’Epilepsie (LFCE). This work has received support from the French government under the “France 2030” program via A*Midex (Initiative d’Excellence d’Aix-Marseille Université, AMX-19-IET-004) and ANR funding (ANR-17-EURE-002).

## Declaration of interests

The authors declare no competing interests.

## Notes

### Competing Interest Statement

The authors have declared no competing interest.

### Summary of Updates

This version includes a update of data and new analysis (particularly for the probe trial during BCM, where we better took into consideration time period during which mice did not move from the release zone).

## Reference List

Amilhon, B., Huh, C.Y.L., Manseau, F., Ducharme, G., Nichol, H., Adamantidis, A., Williams, S., 2015. Parvalbumin Interneurons of Hippocampus Tune Population Activity at Theta Frequency. Neuron 86, 1277–1289. 10.1016/j.neuron.2015.05.027

Ang, C.W., Carlson, G.C., Coulter, D.A., 2006. Massive and Specific Dysregulation of Direct Cortical Input to the Hippocampus in Temporal Lobe Epilepsy. J. Neurosci. 26, 11850–11856. 10.1523/JNEUROSCI.2354-06.2006

Bershteyn, M., Bröer, S., Parekh, M., Maury, Y., Havlicek, S., Kriks, S., Fuentealba, L., Lee, S., Zhou, R., Subramanyam, G., Sezan, M., Sevilla, E.S., Blankenberger, W., Spatazza, J., Zhou, L., Nethercott, H., Traver, D., Hampel, P., Kim, H., Watson, M., Salter, N., Nesterova, A., Au, W., Kriegstein, A., Alvarez-Buylla, A., Rubenstein, J., Banik, G., Bulfone, A., Priest, C., Nicholas, C.R., 2023. Human pallial MGE-type GABAergic interneuron cell therapy for chronic focal epilepsy. Cell Stem Cell 30, 1331–1350.e11. 10.1016/j.stem.2023.08.013

Bocchio, M., Vorobyev, A., Sadeh, S., Brustlein, S., Dard, R., Reichinnek, S., Emiliani, V., Baude, A., Clopath, C., Cossart, R., 2024. Functional networks of inhibitory neurons orchestrate synchrony in the hippocampus. PLOS Biology 22, e3002837. 10.1371/journal.pbio.3002837

Boileau, C., Deforges, S., Peret, A., Scavarda, D., Bartolomei, F., Giles, A., Partouche, N., Gautron, J., Viotti, J., Janowitz, H., Penchet, G., Marchal, C., Lagarde, S., Trebuchon, A., Villeneuve, N., Rumi, J., Marissal, T., Khazipov, R., Khalilov, I., Martineau, F., Maréchal, M., Lepine, A., Milh, M., Figarella-Branger, D., Dougy, E., Tong, S., Appay, R., Baudouin, S., Mercer, A., Smith, J.B., Danos, O., Porter, R., Mulle, C., Crépel, V., 2023. GluK2 Is a Target for Gene Therapy in Drug-Resistant Temporal Lobe Epilepsy. Annals of Neurology 94, 745–761. 10.1002/ana.26723

Buckmaster, P.S., Reyes, B., Kahn, T., Wyeth, M., 2022. Ventral Hippocampal Formation Is the Primary Epileptogenic Zone in a Rat Model of Temporal Lobe Epilepsy. J. Neurosci. 42, 7482–7495. 10.1523/JNEUROSCI.0429-22.2022

Chauvière, L., Rafrafi, N., Thinus-Blanc, C., Bartolomei, F., Esclapez, M., Bernard, C., 2009. Early Deficits in Spatial Memory and Theta Rhythm in Experimental Temporal Lobe Epilepsy. J. Neurosci. 29, 5402–5410. 10.1523/JNEUROSCI.4699-08.2009

Cossart, R., Dinocourt, C., Hirsch, J.C., Merchan-Perez, A., De Felipe, J., Ben-Ari, Y., Esclapez, M., Bernard, C., 2001. Dendritic but not somatic GABAergic inhibition is decreased in experimental epilepsy. Nature Neuroscience 4, 52–62. 10.1038/82900

Costa, B.S., Santos, M.C.V., Rosa, D.V., Schutze, M., Miranda, D.M., Romano-Silva, M.A., 2019. Automated evaluation of hippocampal subfields volumes in mesial temporal lobe epilepsy and its relationship to the surgical outcome. Epilepsy Research 154, 152–156. 10.1016/j.eplepsyres.2019.05.011

Dimidschstein, J., Chen, Q., Tremblay, R., Rogers, S.L., Saldi, G.-A., Guo, L., Xu, Q., Liu, R., Lu, C., Chu, J., Grimley, J.S., Krostag, A.-R., Kaykas, A., Avery, M.C., Rashid, M.S., Baek, M., Jacob, A.L., Smith, G.B., Wilson, D.E., Kosche, G., Kruglikov, I., Rusielewicz, T., Kotak, V.C., Mowery, T.M., Anderson, S.A., Callaway, E.M., Dasen, J.S., Fitzpatrick, D., Fossati, V., Long, M.A., Noggle, S., Reynolds, J.H., Sanes, D.H., Rudy, B., Feng, G., Fishell, G., 2016. A viral strategy for targeting and manipulating interneurons across vertebrate species. Nature Neuroscience 19, 1743–1749. 10.1038/nn.4430

Dinocourt, C., Petanjek, Z., Freund, T.F., BenLJAri, Y., Esclapez, M., 2003. Loss of interneurons innervating pyramidal cell dendrites and axon initial segments in the CA1 region of the hippocampus following pilocarpineLJinduced seizures. Journal of Comparative Neurology 459, 407–425. 10.1002/cne.10622

Donato, F., Rompani, S.B., Caroni, P., 2013. Parvalbumin-expressing basket-cell network plasticity induced by experience regulates adult learning. Nature 504, 272–276. 10.1038/nature12866

Dudok, B., Klein, P.M., Hwaun, E., Lee, B.R., Yao, Z., Fong, O., Bowler, J.C., Terada, S., Sparks, F.T., Szabo, G.G., Farrell, J.S., Berg, J., Daigle, T.L., Tasic, B., Dimidschstein, J., Fishell, G., Losonczy, A., Zeng, H., Soltesz, I., 2021. Alternating sources of perisomatic inhibition during behavior. Neuron 109, 997–1012.e9. 10.1016/j.neuron.2021.01.003

Dudok, B., Klein, P.M., Soltesz, I., 2022. Toward Understanding the Diverse Roles of Perisomatic Interneurons in Epilepsy. Epilepsy Curr 22, 54–60. 10.1177/15357597211053687

Dugladze, T., Vida, I., Tort, A.B., Gross, A., Otahal, J., Heinemann, U., Kopell, N.J., Gloveli, T., 2007. Impaired hippocampal rhythmogenesis in a mouse model of mesial temporal lobe epilepsy. PNAS 104, 17530–17535. 10.1073/pnas.0708301104

Fisahn, A., Pike, F.G., Buhl, E.H., Paulsen, O., 1998. Cholinergic induction of network oscillations at 40 Hz in the hippocampus in vitro. Nature 394, 186–189. 10.1038/28179

Goirand-Lopez, L., Moulinier, M., Vigier, A., Boileau, C., Carleton, A., Muldoon, S.F., Marissal, T., Crépel, V., 2023. Kainate receptors modulate the microstructure of synchrony during dentate gyrus epileptiform activity. Neurobiology of Disease 185, 106260. 10.1016/j.nbd.2023.106260

Guerrero, D.K.R., Balueva, K., Barayeu, U., Baracskay, P., Gridchyn, I., Nardin, M., Roth, C.N., Wulff, P., Csicsvari, J., 2024. Hippocampal cholecystokinin-expressing interneurons regulate temporal coding and contextual learning. Neuron 112, 2045–2061.e10. 10.1016/j.neuron.2024.03.019

Holmes, G.L., 2013. EEG abnormalities as a biomarker for cognitive comorbidities in pharmacoresistant epilepsy. Epilepsia 54, 60–62. 10.1111/epi.12186

Kang, Y.-J., Clement, E.M., Park, I.-H., Greenfield, L.J., Smith, B.N., Lee, S.-H., 2021. Vulnerability of cholecystokinin-expressing GABAergic interneurons in the unilateral intrahippocampal kainate mouse model of temporal lobe epilepsy. Exp Neurol 342, 113724. 10.1016/j.expneurol.2021.113724

Kim, H.K., Gschwind, T., Nguyen, T.M., Bui, A.D., Felong, S., Ampig, K., Suh, D., Ciernia, A.V., Wood, M.A., Soltesz, I., 2020. Optogenetic intervention of seizures improves spatial memory in a mouse model of chronic temporal lobe epilepsy. Epilepsia. 10.1111/epi.16445

Klausberger, T., Somogyi, P., 2008. Neuronal Diversity and Temporal Dynamics: The Unity of Hippocampal Circuit Operations. Science 321, 53–57. 10.1126/science.1149381

Krook-Magnuson, E., Armstrong, C., Oijala, M., Soltesz, I., 2013. On-demand optogenetic control of spontaneous seizures in temporal lobe epilepsy. Nature Communications 4. 10.1038/ncomms2376

Ledri, M., Madsen, M.G., Nikitidou, L., Kirik, D., Kokaia, M., 2014. Global Optogenetic Activation of Inhibitory Interneurons during Epileptiform Activity. J. Neurosci. 34, 3364–3377. 10.1523/JNEUROSCI.2734-13.2014

Lee, S.-H., Kang, Y.-J., Smith, B.N., 2024. Activation of hypoactive parvalbumin-positive fast-spiking interneurons restores dentate inhibition to reduce electrographic seizures in the mouse intrahippocampal kainate model of temporal lobe epilepsy. Neurobiology of Disease 203, 106737. 10.1016/j.nbd.2024.106737

Lehner, A., Hoffmann, L., Rampp, S., Coras, R., Paulsen, F., Frischknecht, R., Hamer, H., Walther, K., Brandner, S., Hofer, W., Pieper, T., Reisch, L.-M., Bien, C.G., Blumcke, I., 2024. Age-dependent increase of perineuronal nets in the human hippocampus and precocious aging in epilepsy. Epilepsia Open n/a. 10.1002/epi4.12963

Lenck-Santini, P.-P., Holmes, G.L., 2008. Altered Phase Precession and Compression of Temporal Sequences by Place Cells in Epileptic Rats. J. Neurosci. 28, 5053–5062. 10.1523/JNEUROSCI.5024-07.2008

Lentini, C., d’Orange, M., Marichal, N., Trottmann, M.-M., Vignoles, R., Foucault, L., Verrier, C., Massera, C., Raineteau, O., Conzelmann, K.-K., Rival-Gervier, S., Depaulis, A., Berninger, B., Heinrich, C., 2021. Reprogramming reactive glia into interneurons reduces chronic seizure activity in a mouse model of mesial temporal lobe epilepsy. Cell Stem Cell 28, 2104–2121.e10. 10.1016/j.stem.2021.09.002

Lévesque, M., Chen, L.-Y., Etter, G., Shiri, Z., Wang, S., Williams, S., Avoli, M., 2019. Paradoxical effects of optogenetic stimulation in mesial temporal lobe epilepsy. Annals of Neurology 86, 714–728. 10.1002/ana.25572

Li, P., Geng, X., Jiang, H., Caccavano, A., Vicini, S., Wu, J., 2019. Measuring Sharp Waves and Oscillatory Population Activity With the Genetically Encoded Calcium Indicator GCaMP6f. Frontiers in Cellular Neuroscience 13.

Ling, Q., Herstine, J.A., Bradbury, A., Gray, S.J., 2023. AAV-based in vivo gene therapy for neurological disorders. Nat Rev Drug Discov 22, 789–806. 10.1038/s41573-023-00766-7

Liu, Y.-Q., Yu, F., Liu, W.-H., He, X.-H., Peng, B.-W., 2014. Dysfunction of hippocampal interneurons in epilepsy. Neurosci. Bull. 30, 985–998. 10.1007/s12264-014-1478-4

Marissal, T., 2021. An inventory of basic research in temporal lobe epilepsy. Revue Neurologique.

Marissal, T., Salazar, R.F., Bertollini, C., Mutel, S., De Roo, M., Rodriguez, I., Müller, D., Carleton, A., 2018. Restoring wild-type-like CA1 network dynamics and behavior during adulthood in a mouse model of schizophrenia. Nature Neuroscience 21, 1412–1420. 10.1038/s41593-018-0225-y

Miri, M.L., Vinck, M., Pant, R., Cardin, J.A., 2018. Altered hippocampal interneuron activity precedes ictal onset. eLife 7, e40750. 10.7554/eLife.40750

Muldoon, S.F., Villette, V., Tressard, T., Malvache, A., Reichinnek, S., Bartolomei, F., Cossart, R., 2015. GABAergic inhibition shapes interictal dynamics in awake epileptic mice. Brain 138, 2875–2890. 10.1093/brain/awv227

Parrish, R.R., Codadu, N.K., Scott, C.M.-G., Trevelyan, A.J., 2019. Feedforward inhibition ahead of ictal wavefronts is provided by both parvalbumin- and somatostatin-expressing interneurons. The Journal of Physiology 597, 2297–2314. 10.1113/JP277749

Peng, Z., Zhang, N., Wei, W., Huang, C.S., Cetina, Y., Otis, T.S., Houser, C.R., 2013. A Reorganized GABAergic Circuit in a Model of Epilepsy: Evidence from Optogenetic Labeling and Stimulation of Somatostatin Interneurons. J. Neurosci. 33, 14392–14405. 10.1523/JNEUROSCI.2045-13.2013

Peret, A., Christie, L.A., Ouedraogo, D.W., Gorlewicz, A., Epsztein, J., Mulle, C., Crépel, V., 2014. Contribution of Aberrant GluK2-Containing Kainate Receptors to Chronic Seizures in Temporal Lobe Epilepsy. Cell Reports 8, 347–354. 10.1016/j.celrep.2014.06.032

Proddutur, A., Nguyen, S., Yeh, C.-W., Gupta, A., Santhakumar, V., 2023. RECLUSIVE CHANDELIERS: FUNCTIONAL ISOLATION OF DENTATE AXO-AXONIC CELLS AFTER EXPERIMENTAL STATUS EPILEPTICUS. Prog Neurobiol 231, 102542. 10.1016/j.pneurobio.2023.102542

Shuman, T., Aharoni, D., Cai, D.J., Lee, C.R., Chavlis, S., Page-Harley, L., Vetere, L.M., Feng, Y., Yang, C.Y., Mollinedo-Gajate, I., Chen, L., Pennington, Z.T., Taxidis, J., Flores, S.E., Cheng, K., Javaherian, M., Kaba, C.C., Rao, N., La-Vu, M., Pandi, I., Shtrahman, M., Bakhurin, K.I., Masmanidis, S.C., Khakh, B.S., Poirazi, P., Silva, A.J., Golshani, P., 2020. Breakdown of spatial coding and interneuron synchronization in epileptic mice. Nat Neurosci 1–10. 10.1038/s41593-019-0559-0

Toyoda, I., Bower, M.R., Leyva, F., Buckmaster, P.S., 2013. Early activation of ventral hippocampus and subiculum during spontaneous seizures in a rat model of temporal lobe epilepsy. J Neurosci 33, 11100–11115. 10.1523/JNEUROSCI.0472-13.2013

Vigier, A., Partouche, N., Michel, F.J., Crépel, V., Marissal, T., 2021. Substantial outcome improvement using a refined pilocarpine mouse model of temporal lobe epilepsy. Neurobiology of Disease 161, 105547. 10.1016/j.nbd.2021.105547

Wang, Ying, Liang, J., Chen, L., Shen, Y., Zhao, J., Xu, C., Wu, X., Cheng, H., Ying, X., Guo, Y., Wang, S., Zhou, Y., Wang, Yi, Chen, Z., 2018. Pharmaco-genetic therapeutics targeting parvalbumin neurons attenuate temporal lobe epilepsy. Neurobiology of Disease 117, 149–160. 10.1016/j.nbd.2018.06.006

Wang, Ying, Wang, Yi, Xu, C., Wang, S., Tan, N., Chen, C., Chen, L., Wu, X., Fei, F., Cheng, H., Lin, W., Qi, Y., Chen, B., Liang, J., Zhao, J., Xu, Z., Guo, Y., Zhang, S., Li, X., Zhou, Y., Duan, S., Chen, Z., 2020. Direct Septum-Hippocampus Cholinergic Circuit Attenuates Seizure Through Driving Somatostatin Inhibition. Biological Psychiatry, Neurodegeneration 87, 843–856. 10.1016/j.biopsych.2019.11.014

Wozny, C., Gabriel, S., Jandova, K., Schulze, K., Heinemann, U., Behr, J., 2005. Entorhinal cortex entrains epileptiform activity in CA1 in pilocarpine-treated rats. Neurobiology of Disease 19, 451–460. 10.1016/j.nbd.2005.01.016

Zhan, J., Komal, R., Keenan, W.T., Hattar, S., Fernandez, D.C., 2019. Non-invasive Strategies for Chronic Manipulation of DREADD-controlled Neuronal Activity. J Vis Exp 10.3791/59439. 10.3791/59439

