## Supplemental material for "Minimally-invasive Manipulation of Spared and Hypoactive Interneurons reduces CA1 Synchronization and Nonspatial Behavior alterations in Epilepsy models"

1. **Supplemental Materials and Methods:**

**Animals**

We used wild-type mice from Swiss/CD-1 background (from Charles River and Janvier Labs). Animals were kept on a normal 12h light/dark cycle at 22–24°C and with ad libitum access to food and water. Control and epileptic mice were single housed in enriched cages. All experiments conducted on mice were performed in accordance with the directives of European Community Council (2010/63/UE) and received approval from the French Ministry for Research, after ethical evaluation by the institutional animal care and use committee of Aix-Marseille University (APAFIS number: #23627). Across all experiments, particular care was taken to minimize any stress or suffering in animals, including manipulating them using a tunnel to avoid grabbing them by the tail.

**In vivo pilocarpine-based model of TLE**

To examine the contribution of interneurons in cognitive comorbidities *in vivo*, we used a low-mortality pilocarpine-based procedure to generate mice that exhibit histological and electrophysiological characteristics associated with TLE (Vigier et al., 2021). To avoid the difference due to circadian effects, mice underwent the pilocarpine TLE induction at the same time of the day (i.e., starting from around 10:00 AM). In brief, we placed adult males (2-3 months old) in individual cylindrical tanks made from transparent Plexiglas (diameter: 30 cm, height: 40 cm). Mice were administered with an intraperitoneal injection of scopolamine methyl-nitrate (10 mg/kg, IP, ChemCruz) to reduce the peripheral cholinergic effects of pilocarpine. Thirty minutes later, we applied a protocol with a ramp-up pilocarpine dose until the development of *Status Epilepticus* (SE). A first dose of pilocarpine was administered (300 mg/kg, IP, Sigma) and the mouse behavior was monitored. As long as the mice did not develop short seizure with motor manifestation (myoclonus), we injected 150 mg/kg of pilocarpine every 30 minutes. If mice experienced myoclonus, we performed a single injection of caffeine (40 mg/kg, IP, Sigma). SE generally occurred within the following 60 min. After 1 hour of SE, the mice were injected with diazepam to interrupt seizure activity (10 mg/kg, IP, Laboratoire TVM). After the protocol, the animals were injected with a saline solution to avoid dehydration (0.9%, Osalia, subcutaneous). They were then placed in individual cages containing enrichment and a hyper-nourishing gel (Gel-Safe). The mice were observed daily for 3 weeks, then at least twice a week until death.

**In vitro pilocarpine-based model of TLE**

To investigate the relationship between the recruitment of interneurons and the CA1 epileptic network dynamics *in vitro* as in (Cǎlin et al., 2018), we used long lifespan mouse hippocampal organotypic slices (referred to as “slices” for simplicity), which display, after application of pilocarpine, histological and network dynamics alterations that are similar to those observed in the mouse *in vivo* pilocarpine-based model of TLE that we use, as well as tissues from patients with TLE (Boileau et al., 2023; Goirand-Lopez et al., 2023; Peret et al., 2014). 5–7-days-old mice of both sexes were decapitated, and their hippocampi were removed and unfolded in PBS + 30mM Glucose. Transverse hippocampal slices (400 μm thickness) were made using a chopper (McIlwain). Slices were placed on Millicell-CM Biopore membranes (Millipore) in culture dishes containing 1 ml of the following medium: MEM 50%, HS 25%, HBSS 25%, HEPES 15 mM, glucose 6.5 mg/ml, and insulin 0.1 mg/ml. Slices were maintained in an incubator at 37 ◦C/5% CO2, and the culture medium was changed every 2–3 days. Viral infection of the slices was performed the day after the preparation of the culture (i.e., day *in vitro* 1, DIV1), by applying approximately 300 nl of adeno-associated-virus- (AAV-)containing solution onto the surface of the hippocampus. To induce the histological damages and network reorganization similar to those observed in mouse TLE models and patients (Boileau et al., 2023; Goirand-Lopez et al., 2023; Peret et al., 2014), 0.5 μM of pilocarpine was added to the culture medium from *in vitro* day 3 to 6 (DIV3-6). Calcium imaging experiments were conducted a minimum of one week after the pilocarpine treatment, i.e., from DIV10 to DIV14.

**Surgeries**

Surgery was performed in a subgroup of mice to inject AAVs enabling the manipulation of inhibitory neurons and/or to record ictal activity. In both cases, mice were subcutaneously injected with an analgesic (Buprenorphine, 45μg/kg) and an anti-inflammatory drug (Carprofene, 10mg/kg) 20-30 minutes prior to the surgery. Anesthesia was induced at 5% in a transparent chamber (Minerve) and maintained at 0.5–3% isoflurane (w/v) using a mouse gas mask (Minerve) in a stereotaxic apparatus (Stoelting Co.) coupled to a binocular microscope (SZ61, Olympus). Mice were maintained at 36.5–38**°**C using a heating pad and controlled via a rectal thermometer (Stoelting). Mouse heads were secured using ear bars, a jaw bar, and a nose clamp. Eyes were covered with ointment (Ocry-Gel). Lurocaïne (Lidocaine, 7mg/kg), a local analgesic, was injected below the skin overlaying the skull. Then, the two procedures differed:

***Viral injections***

AAVs were injected bilaterally in the dorsal and ventral hippocampus of chronically epileptic animals minimum 3 weeks after TLE induction or age-matched controls. We used AAV constructs enabling the expression of the excitatory chemogenetic “DREADD” receptor hM3D and the red reporter tdTomato under the control of the interneuron-specific hDlx enhancer element (pAAV-hDlx-GqDREADD-tdTomato-Fishell-4, AAV9, titer > 7×10^12^ vg/mL; these AAVs were a gift from Gordon Fishell, Addgene, 162375-AAV9) (Dimidschstein et al., 2016). The coordinates for the dorsal and ventral hippocampi were: AP -1.8, ML ± 1, DV -2 and AP -3.3, ML ±2.3, DV -2.5 from the bregma, respectively. In brief, the scalp of the anesthetized animals was shaved using electric clippers and then incised using a scalpel blade. The skull bone was gently cleaned and dried. The position of the craniotomies was then defined with a fine marker. The skull was pierced using a dental driller (Fordedim), without damaging the meninges. Then, 500nL of AAV was injected using a Nanoject III injector (Drummond) and glass capillaries (rate: 20nL/seconds, 10 cycles separated by 1 second). After injections, the skin was sutured with surgical stitches (Ethilon 5-0 polyamide thread), then anesthesia was discontinued. This protocol enables robust expression of genes of interest in the hippocampus, particularly in CA1. Expression of genes of interest has also been reported in the cortex above the hippocampus, which is traversed by the injection capillary.

***Implantation of the EEG recording device***

A minimum of two weeks after AAV injection, the scalp and the back of the animals were shaved using electric clippers. The skin was cleaned with antiseptic (Vetadine) and sterile saline solution (0.9%, Osalia). The skull was exposed using a scalpel blade. Three craniotomies were made using a drill (Foredom): two in an intermediate position between the two AAV injection sites in both hemispheres (approximately AP -2.55 mm, ML +1.65 mm relative to Bregma), the last above the cerebellum as in (Boileau et al., 2023). Then, the mouse back was incised, and a subcutaneous pouch was created by blunt dissection. A radio-frequency transmitter (ETA-F10, DSI) was implanted inside the pocket close to the ventral abdominal region, under the left flank, with wires oriented towards the head. The two emitter wires were carefully slipped underneath the mouse’s neck skin. The tip of the first wire was inserted into the hole above the right hippocampus (~1.5mm from skull surface). The second wire, placed above the cerebellum (~2 mm from the skull surface), served as the ground electrode. To fix the wires, a screw was put in each of the three holes and covered with dental cement (Paladur). Finally, the cranial and flank incisions were sutured using 5-0 polyamide filaments (Ethilon, Ethicon). Mice were injected with saline solution (0.9%, 0.35mL) and allowed to recover from the surgery on a heating pad until fully awakened. Animal’s weight was then monitored. Buprenorphine (45µg/kg) was injected 24 hours after surgery, and Carprofen (10mg/kg) and saline solution (0.9%, 0.35mL) were injected 24 and 48 hours after surgery, all subcutaneously.

**Behavioral testing**

We performed various tests to measure mouse behavior in different experimental conditions: pilocarpine-treated chronically epileptic animals two months after TLE induction (referred to as “Pilo”) and age-matched controls, injected with AAVs, with normal drinking water, drinking water with 4% of sucrose (a condition referred to as “vehicle” or “VEH”), or clozapine-N-oxide (CNO, Tocris) diluted in the 4%-sucrose drinking water (5 mg/200 mL), referred as “CNO”. Animal assignment to the various experimental groups was randomized. The administration of CNO or VEH started one week before the behavior test (i.e., 1-2 weeks after viral injection) and ended at its completion. The administration of CNO in water is a less invasive method than daily intraperitoneal injections (Zhan et al., 2019). The sucrose was used to cover the taste of CNO, which is postulated to be slightly bitter (Milosavljevic et al., 2016). Drinking water consumption was measured regularly to calculate the average CNO intake per animal per day. Particular care was taken when manipulating the bottles to limit waste of treated water. Mice were handled by the experimenter every day for a week and habituated to the experimental room the day before behavioral testing. Tests were performed under indirect room lighting conditions (120 lux) and at similar times of the day difference due to circadian effects (i.e., starting from 10:00AM). Mouse behavior was continuously videotaped by a video camera placed above the apparatus, tracked using the Ethovision 11.5 software, and the detection was manually corrected if necessary. The apparatuses were cleaned with 30% ethanol before each trial to eliminate olfactory cues. When a mouse experienced a seizure with motor manifestation before or during the behavioral testing, it was returned to its home cage, then tested again at least 30 minutes later. However, in the absence of EEG monitoring during behavior, we cannot exclude that epileptiform activities without obvious motor signs occurred during the experiment. All criteria used for including and excluding animals were defined prior to analysis. Across all experiments, we discarded any mouse that presented ulcerative dermatitis, a skin affection that can be observed in pilocarpine-treated animals after repetitive scratching. We also excluded from analysis animals that presented abnormal locomotion, i.e., stereotyped movements, or inability to move normally. For exclusion criteria in specific tests, see below for details.

***Barnes circular maze (BCM)***

BCM (Gawel et al., 2019) consisted of an elevated circular platform made of metal (diameter = 100 cm; elevation from the ground = 90 cm, color: grey) containing 20 holes equally spaced around the periphery (diameter = 4 cm)(**Fig. 4C**). Under one hole was placed a magnetic escape room (black) in which mice could hide. Under the remaining 19 holes were placed black caps, which, from the surface of the platform, appeared indistinguishable from the escape room. Distal visual cues (length/width = 30 cm) surrounded the platform. BCM was performed while a loudspeaker played white noise (50-60dB). 24h prior to the test, mice were allowed to freely explore the maze during 1-2 habituation sessions of 3 minutes after which they were gently guided to the escape room. Then, mice underwent the acquisition phase of the test (during which spatial learning-related abilities were assessed). This acquisition phase consisted of 2 trials per day for 4 consecutive days during which the position of the escape room remained constant. For each trial, mice were placed in a start chamber in the middle of the maze (“release zone”). After 15 seconds, the chamber was lifted, and the animals were allowed to explore the maze. The trial concluded when the mice entered the escape room, or after 2 minutes elapsed. If a mouse failed to enter the escape room within the 2-minute period, it was gently guided to the escape room by the operator and left there for 15 seconds prior to removal. For each day of habituation and acquisition, several parameters were quantified to assess mouse performance, including: the success and failure rate (i.e., the % of mice per experimental group that identified or not the escape room location at least once per day of acquisition), the latency to locate the escape room (“primary latency”), along with the number of incorrect holes checked prior to finding the escape room or the completion of the test (“primary errors”). Some pilocarpine-treated animals jumped off the apparatus and were considered to have failed the test session. To ensure correct learning, they were quickly returned to the table to complete the test under the same conditions as the other animals. For quantification of primary latencies and errors, we calculated the average of the non-null values for each trial per day of the BCM acquisition phase. We also classified the mouse exploration strategy as direct (when mice explored less than 3 distinct holes prior to finding the escape room), serial (when mice explored consecutive holes separated by up to 2 holes in a clockwise or counterclockwise manner) or random (when mice crossed the maze center, ran circularly on the periphery with no or few hole inspections, used mixed strategies, jumped off the apparatus or poorly explored the apparatus during the duration of a trial, i.e., less than 5 distinct holes explored) (Gawel et al., 2019). To assess behaviors related to spatial memory, animals underwent the probe trial 72 hours after the last acquisition day. During the probe trial, the escape room was removed and replaced by a false escape box (identical to the cap placed underneath the other 19 holes). The probe trial was performed in similarly to the acquisition trials (15 seconds in the start chamber, 2 minutes of free exploration). The success rate, the latency prior to reaching to previous escape room location, the number of incorrect holes checked, and exploration strategies were measured, along with the time spent in the quadrant where the escape room was previously localized (“target zone”). We measured the distance covered by the mice (giving an indication of locomotory activity) only during the BCM probe trial, given that it is the only stage of the test during which the animals remained on the apparatus for the same amount of time, except when epileptic mice jumped off the apparatus (which we reported in **Supplementary Fig. 4A**). Given the extent and variability in the altered BCM-related behavior of epileptic mice, and in order to avoid missing or aberrant values, we performed a min-max normalization for latency and number of primary errors, as well as time in the target area, a score of 0 being assigned to the best possible performance (e.g., the mouse quickly finds the escape room without making any error during the acquisition phase, or moves into the target zone during the whole probe trial) and 1 to the worst possible performance. Thus, a score of 1 was assigned for primary latency if a mouse failed to find the escape room location or jumped off the apparatus before finding the escape room. A score of 1 was assigned for primary errors when an animal jumped off the apparatus before finding the escape room location or failed to locate the escape room location after little exploration of the apparatus, i.e., 0-5 distinct holes explored. A score of 1 was assigned for the time in target area if an animal jumped off the apparatus, or did not move from the release zone for the entire trial period. The original data is shown in **Supplemental Fig. 4A**.

***Novel Object Recognition test (NORT)***

NORT was performed to measure mouse non-spatial memory (Antunes and Biala, 2012) (**Fig. 4F-G and Supplemental Fig. 4B**), which involves the dorsal CA1 (Cohen et al., 2013). NORT was performed in a dark Plexiglas box (40 cm L x 40 cm W x 40 cm H). After habituation to the arena (10 minutes, 1 day prior to the familiarization stage), mice were placed into the center of the box, in which two identical objects were placed in opposite quarters. During this 5-minutes familiarization stage, the time exploring each object was measured. Mice were defined as exploring the object when their nose was directed towards the object at a distance of < 2 cm. Mice that remained for >100 seconds in a quarter of the apparatus without exploratory behavior were excluded from the analysis. 1 day later, during the testing phase, one of the objects was replaced by a new item of similar size, and the time exploring the familiar versus the novel object was measured for 5 minutes. To limit bias due to an animal's preference for a particular location in the apparatus, the new object was systematically replacing the object explored the least in the previous phase. Used objects included brick toys, 0.5L bottles, and needle boxes.

**Histology**

The week following the end of behavioral testing, mice were anesthetized (using a mix of Domitor 1.2mg/kg and Zoletil 80mg/kg) prior to intracardiac perfusion with PBS (ThermoFisher)/Heparin (0.1%, Sigma) solution and then with a paraformaldehyde-based solution (Antigenfix, Diapath). Next, the animals were decapitated, and the brains were kept for a maximum of 24 h in Antigenfix at 4 ◦C before being washed in PBS. Brains were placed overnight in 30% sucrose at 4◦C for cryopreservation. Afterward, brains were embedded in blocks containing optimum cutting temperature medium (OCT; Tissue-Tek) placed on dry ice, enabling tissue storage at 80◦C for months if necessary. Finally, 60-μm-thick coronal sections were prepared at -25°C from the brain tissue blocks using a Cryostat (Leica CM3050S). Sections were stored in 24 well plates according to their position in the anteroposterior axis of the brain at 4°C in PBS-Azide (0.01%) until immunochemistry was performed. We focused our histological study on dorsal CA1. To enable subsequent comparison between conditions, we systematically selected for each animal two sections that are situated at 840 and 1200 µm posterior to the most anterior part of the hippocampus. These sections were permeabilized with PBS-Triton (1%) containing horse serum (10%) for 2 h at room temperature. Next, sections were incubated with polyclonal goat primary antibody directed against Parvalbumin (PV, Swant, 1:1000) and monoclonal mouse primary antibody directed against Somatostatin (SOM, Santa-Cruz, 1:200), or polyclonal guinea pig primary antibody directed against CCK8 (Synaptic System, 1:1000) in PBS-Triton (1%) containing horse serum (10%) for 1-2 days at 4 ◦C under agitation. Sections were then washed with PBS, placed in PBS containing polyclonal donkey secondary antibody coupled with Alexa488 directed against goat (Invitrogen, 1:500) and donkey secondary antibody coupled with Alexa647 directed against mouse (Invitrogen, 1:500), or with donkey secondary antibody coupled with Cyanine Cy5 directed against guinea pig (Jackson ImmunoResearch Laboratories, 1:500) for 2 h at room temperature under agitation. Sections were then washed with PBS, and stained with Hoechst (Sigma, 1:2000) at room temperature under agitation for 5-10 min before mounting (Fluoromount-G mounting medium, Invitrogen). 4 images per animal (2 in each hemisphere) were acquired using a confocal microscope (LSM800, Zeiss equipped with 10x/0.3 and 20×/0.8 objectives). Immunofluorescence was detected in a field of X638.9 × Y638.9 μm (Z-stacks of 8 planes, interval 2 µm) covering all sublayers of dorsal CA1 from the alveus to the stratum lacunosum-moleculare (**Fig. 1B**). Images were processed and quantification was performed manually using ImageJ software. In brief, the area of quantified CA1 was measured and the number of PV, SOM, Cholecystokinin (CCK) and tdTomato-expressing interneurons per animal was counted in the 4 images on z-stacks plane by plane and not on projections. Then, density per animal was estimated by dividing the number of somata by the volume of quantified CA1, averaged over the 4 images. Additionally, in some control and epileptic animals, low magnification mosaic images of the Hoechst-stained hippocampi (X6172 × Y2695 μm) were acquired to measure CA1 area (**Supplemental** **Fig. 1A-B**). For interneurons density quantification, images of epileptic animals presenting a massive sclerosis of CA1 (i.e., stratum pyramidale no longer visible) were excluded from the analysis due to difficulties assessing properly the borders of CA1 and its subregions. In all conditions, we also excluded images presenting damaged tissue attributable to histological processing.

**Calcium imaging and analysis**

To monitor the activity of a large population of CA1 neurons, slices were transduced with AAV constructs enabling the expression of the genetically encoded calcium sensor jGCaMP8m under the control of the human synapsin promoter (pGP-AAV-syn-jGCaMP8m-WPRE, AAV9, titer > 1×10^13^ vg/mL). pGP-AAV-syn-jGCaMP8m-WPRE was a gift from GENIE Project (Addgene, 162378-AAV9) (Zhang et al., 2023). To identify and manipulate CA1 interneurons, slices were transduced with AAV9-hDlx-GqDREADD-tdTomato-Fishell-4 (Dimidschstein et al., 2016). On the *in vitro* days 10-14, slices were immersed in an artificial cerebrospinal fluid (ACSF) containing (in mM): 126 NaCl, 3.5 KCl, 1.2 NaH 2PO4, 26 NaHCO 3, 1.3 MgCl2, 2.0 CaCl2, and 10 Glucose (pH ~7.4), continuously oxygenated and perfused at ~30 ◦C using a peristaltic pump. All compounds used for the preparation of the ACSF were purchased from Sigma-Aldrich. Calcium transients were recorded using a multibeam two-photon laser scanning system (TriMScope, LaVision Biotec) mounted on a BX61WI Olympus microscope coupled to an Ultra II chameleon laser (Coherent). A fluorescent lamp (Olympus U-RFL-T) was used to identify a CA1 field with GCaMP-expressing neurons and tdTomato-expressing interneurons. Slices were imaged using a 20 X 0.95NA objective (Olympus) at a ~23 Hz frame rate (i.e., 44 ms per frame) for up to ~7 minutes per recording (i.e., 10,000 frames). The two-photon excitation wavelength was 920 nm. A typical imaging session covered a field of 300 × 150 μm size containing up to 127 individual CA1 neurons. To record the population activity in control versus pathological conditions as in (Li et al., 2019; Marissal et al., 2018), GCaMP-based calcium activity was recorded under the continuous presence of 50µM carbachol (Cch, Sigma). The application of Cch on hippocampal slices *in vitro* notably induces oscillatory activity resembling those that occur *in vivo* during navigation, learning and memory processes (Fisahn et al., 1998; Li et al., 2019; Marissal et al., 2018). To test whether Cch-induced CA1 calcium activity reflects the neuronal action potential discharge, some slices were monitored twice, the first time in Cch only, then the sodium channel blocker tetrodotoxin (TTX, Tocris, 1µM) was applied in the bath in addition to the Cch for at least 15 minutes, followed by a second calcium imaging of the same slice. In another subset of experiments, to test the effect of interneuron manipulation on the network dynamics, some slices were monitored first in Cch only, then the DREADD ligand Clozapine-N-Oxide (CNO, Tocris, 2µM) was applied in the bath in addition to the Cch for at least 15 minutes, followed by a second calcium imaging of the same slice. 15-minute bath application of CNO was shown sufficient to elevate the activity of all interneurons expressing excitatory DREADDs (Marissal et al., 2018). The detection of neurons and calcium activity onsets/offsets was performed semi-automatically in each movie using the Matlab-based ‘Caltracer’ software (Columbia University). Typical calcium events were characterized by a sharp rise and a slow decay, lasting for 1 second and with an amplitude of at least twice the baseline. Then, we calculated the level of calcium activity (i.e., number of active neurons, frequency, amplitude, and duration of calcium transients), and quantified neuronal synchronization as in patients (Goirand-Lopez et al., 2023; Marissal et al., 2018). Calcium imaging recordings in which the slice displayed signs of degradation (e.g., very bright neurons with fluorescence in the nucleus) were excluded from the analysis.

**EEG recordings and analysis**

We performed EEG recordings to measure seizure activity in epileptic mice injected with AAV9-hDlx-GqDREADD-tdTomato-Fishell-4 before and after chemo-activation of hippocampal interneurons using CNO. On average 14 weeks after TLE induction and the week following EEG implantation surgery, individual cages containing implanted animals were placed on telemetric receiver plates, and EEG signals (1000× amplified, filtered at 0.16–97 Hz pass, and acquired at 500 Hz) were monitored 24 hours per day for minimum 7 consecutive days. All mice were first recorded during a baseline period with a vehicle in the drinking water (4% sucrose in the drinking water, ad libitum, for 5 days). Then, a group of animals was administered CNO (5mg in 200 mL of 4% sucrose drinking water, ad libitum, for 5 days). These animals (*n =* 9) were called “Pilo-CNO”. For comparison, a group of animals (*n =* 4) were treated with the vehicle throughout the recording. This group is referred to as “Pilo-VEH”. Day 0 corresponds to when CNO was administered to Pilo-CNO mice (and to the sixth day of recording in Pilo-VEH mice). The long duration of recordings was justified by the previous observation that spontaneous ictal events occur in clusters that can be separated by long seizure-free periods (Lim et al., 2018; Vigier et al., 2021). Drinking water consumption was measured regularly to calculate the average CNO intake per animal per day.

EEG was analyzed using the Neuroscore software (DSI). EEG raw signal was visually inspected and the ‘periodogram’ function of DSI (time-frequency spectrogram of EEG, determined using a Fast Fourier Transform algorithm with a sliding 10 seconds-Hanning window) was used to help detect ictal events. Similar to (Boileau et al., 2023; Vigier et al., 2021), we quantified in each experimental conditions the frequency and duration of ictal activity characterized by rhythmic (4-60Hz) and prolonged (>10s) spike trains with an amplitude exceeding twice the baseline level. To limit the potential effect of anesthesia and post-surgical care (see Surgery section above), we only analyzed EEG signals recorded from the third day after implantation as in (Padmasola et al., 2024). Time periods during which we observed artifacts (e.g., due to repeated head scratching by the animal) were removed from the analysis if they did not represent a prominent part of the recordings. Otherwise, the whole recording was excluded from analysis.

**Statistical methods**

For the comparison of multiple groups of two factors, we used the two-way ANOVA test followed by Sidak's post-hoc test when appropriate. For comparisons between two unpaired groups with normal distribution, the two-tailed Student's t-test was used. For comparisons between two unpaired groups without normal distribution, the two-tailed Mann-Whitney test was used. For comparisons between two paired groups with normal distribution, the two-tailed paired t-test was used. For comparisons between two paired groups without normal distribution, the two-tailed Wilcoxon test was used. Normality was assessed using the Shapiro-Wilk test. For the analysis of categorical outcomes (e.g., success or failure), we used Fisher’s exact test. All statistical tests were performed using GraphPad Prism software and detailed in the figure legends. Sample size was decided using InVivoStat software.

2. **Supplemental Figures:**

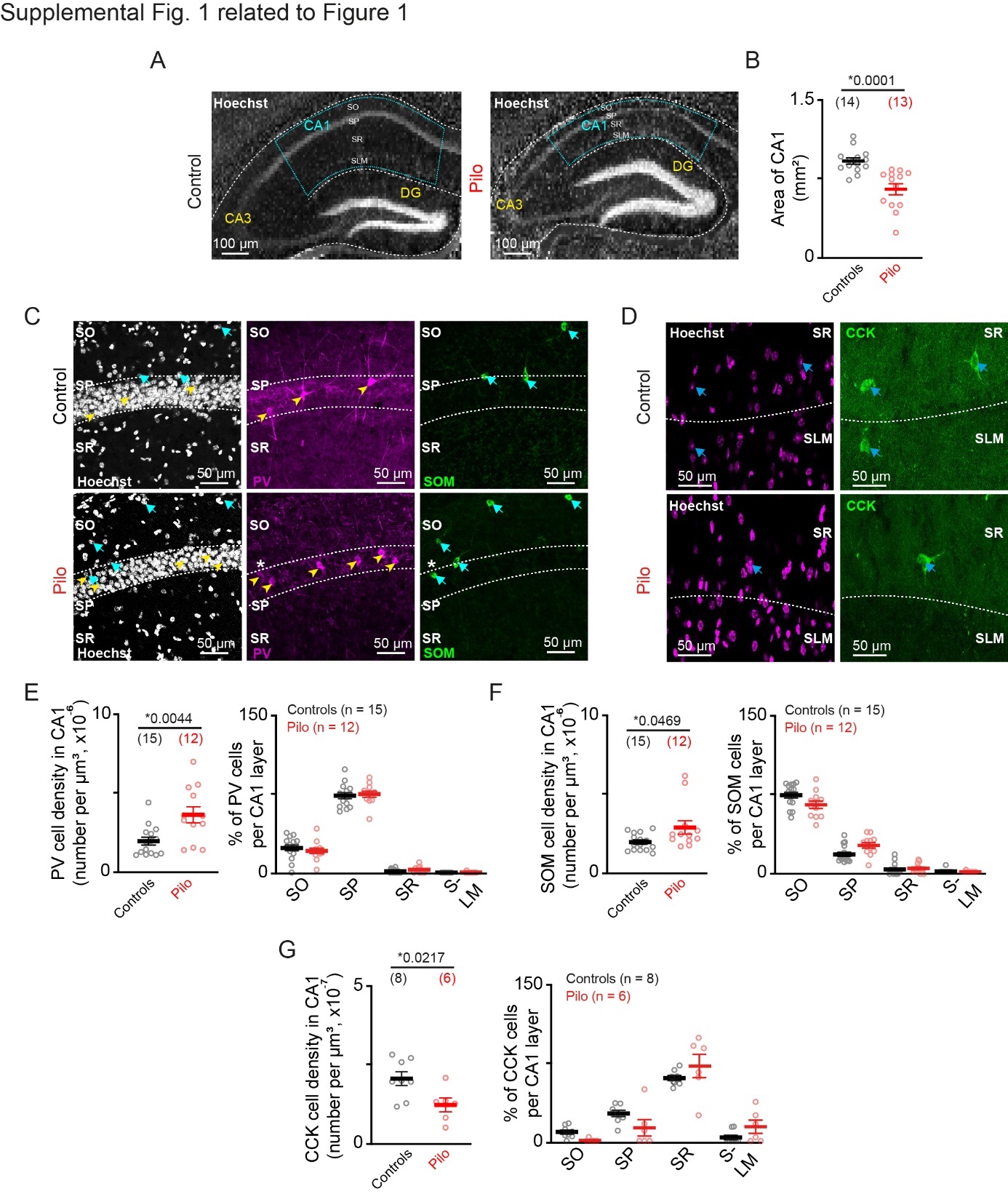

**Supplementary Fig. 1 related to Fig. 1. A.** Representative Hoechst staining in the hippocampus of a Control and a Pilo mouse. In this example from a Pilo mouse, histological alterations have been observed in the hippocampus, such as a reduction in the size of CA1 (delimited with blue dot lines), whose stratum pyramidale (SP) is reduced to a thin layer; as well as dispersion of the granular cells of the dentate gyrus (DG). **B.** Quantification of CA1 area in each condition (Two-tailed Mann-Whitney test). **C.** Representative Hoechst staining (left), as well as Parvalbumin (PV, middle) and Somatostatin (SOM, right) immunolabelling in the CA1 region from a Control (top) or a Pilo (bottom) mouse. Neurons co-labelled with Hoechst and PV or SOM are indicated by a yellow arrowhead or cyan arrow, respectively. A neuron that co-express PV and SOM was indicated by a white asterisk. **D.** Representative Hoechst staining (left), as well as Cholecystokinin (CCK, right) immunolabelling in the CA1 region from a Control (top) or a Pilo (bottom) mouse. Neurons co-labelled with Hoechst and CCK are indicated by a blue arrow. **E-G.** Density (left) and percentage distribution (right) of PV- (E.), SOM- (F.) and CCK-expressing neurons (G.). Density was quantified using Two-tailed Mann-Whitney for all interneuron subtypes. Distribution was quantified using two-way ANOVAs. For PV: two-way ANOVA experimental condition effect F (1, 25) = 0.7937, *P =* 0.3815; for SOM: two-way ANOVA experimental condition effect F (1, 25) = 3.068, *P =* 0.0921); for CCK two-way ANOVA experimental condition effect F (1, 12) = 0.4243, *P =* 0.5271. SO: Stratum Oriens, SP: Stratum Pyramidale, SR: Stratum Radiatum, SLM: Stratum Lacunosum Moleculare. Asterisks indicate significant differences (*P <* 0.05) between the Control and Pilo groups. Data are presented as mean ± s.e.m. Each circle represents a given mouse (The number of mice is specified in parentheses).

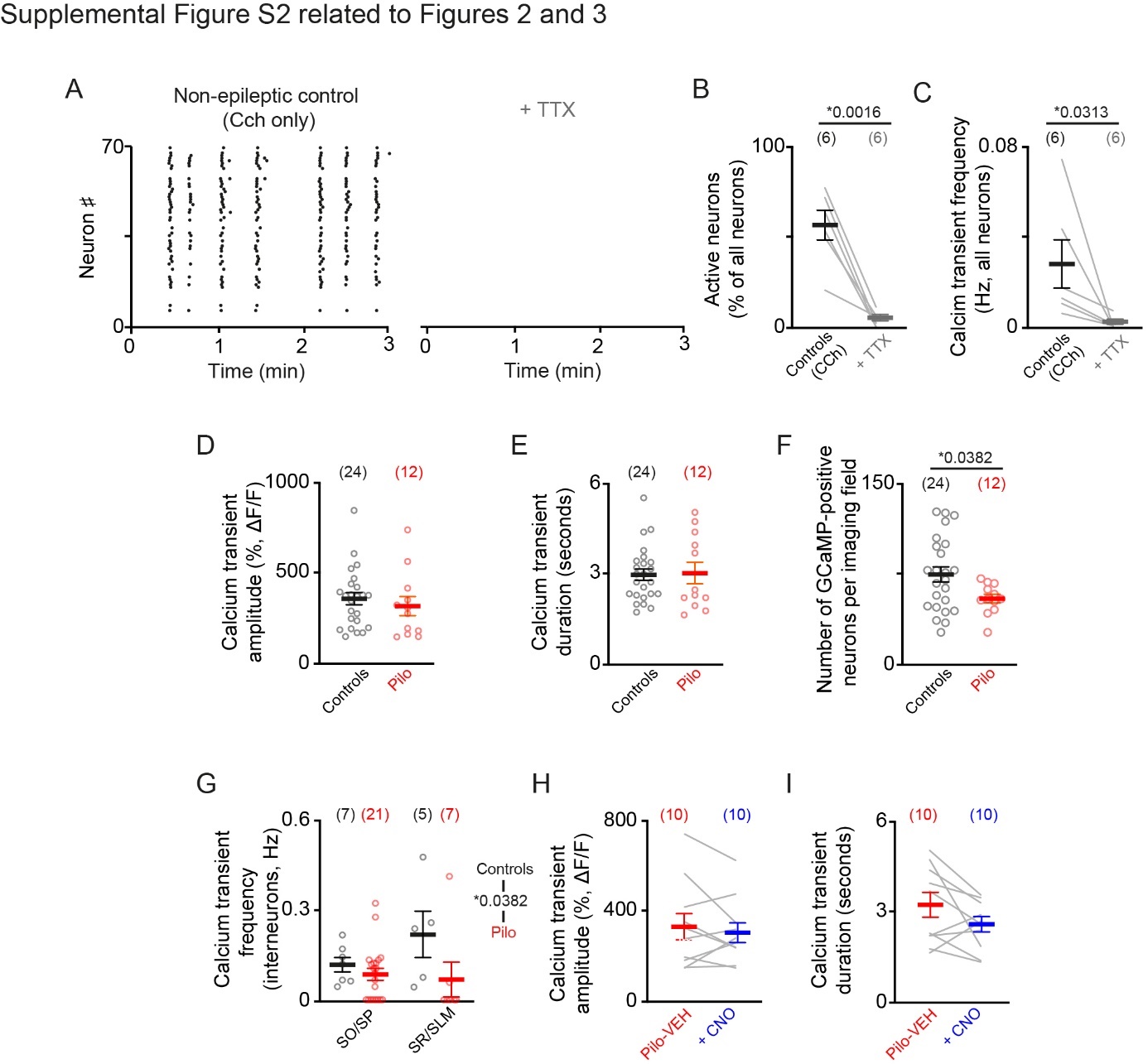

**Supplemental Fig. 2 related to Figs. 2 and 3. A.** Representative raster plots of calcium transients recorded in the continuous presence of carbachol (Cch, 50µM) in control condition, before (black dots, left) and after application of the sodium channel blocker Tetrodotoxin (TTX, grey, right). **B-C.** Proportions of CA1 neurons displaying at least one calcium transient during the duration of the movie (B), and frequency of these calcium transients (C) in control (black) or TTX (grey) conditions. B: Two-tailed paired t-test, C: Wilcoxon test. **D-E.** Calcium transient amplitude (D) and duration (E) in control or pilocarpine-treated (Pilo) slices (Two-tailed Mann-Whitney test, *P =* 0.2760 for amplitude and 0.7721 for duration) in each condition. **F.** Number of GCaMP-expressing neurons per calcium imaging field (Two-tailed unpaired t-test). **G.** Frequency of calcium transients recorded in individual CA1 interneurons localized in deep CA1 layers (i.e., strata oriens and pyramidale, SO/SP) and superficial CA1 layers (i.e., strata radiatum and lacunosum moleculare, SR/SLM) in each condition (Two-way repeated measures ANOVA: experimental group effect F (1, 36) = 4.998, Two-way repeated measures ANOVA location effect F (1, 36) = 1.031, *P =* 0.3167). **H-I.** Calcium transient amplitude (H) and duration (I) in pilocarpine-treated slices before (pilo-VEH) and after treatment with CNO to activate hM3D-expressing interneurons (Two-tailed paired t-test, *P =* 0.4536 and 0.1053, respectively). Data are presented as mean ± s.e.m. Each circle or line represents a given slice (The number of slices is specified in parentheses). A-C: Slices were prepared from *N =* 3 animals in control condition. D-G: Slices were prepared from *N =* 7 and 5 animals in control and Pilo conditions, respectively. H-I: Slices were prepared from *N =* 3 animals in Pilo condition.

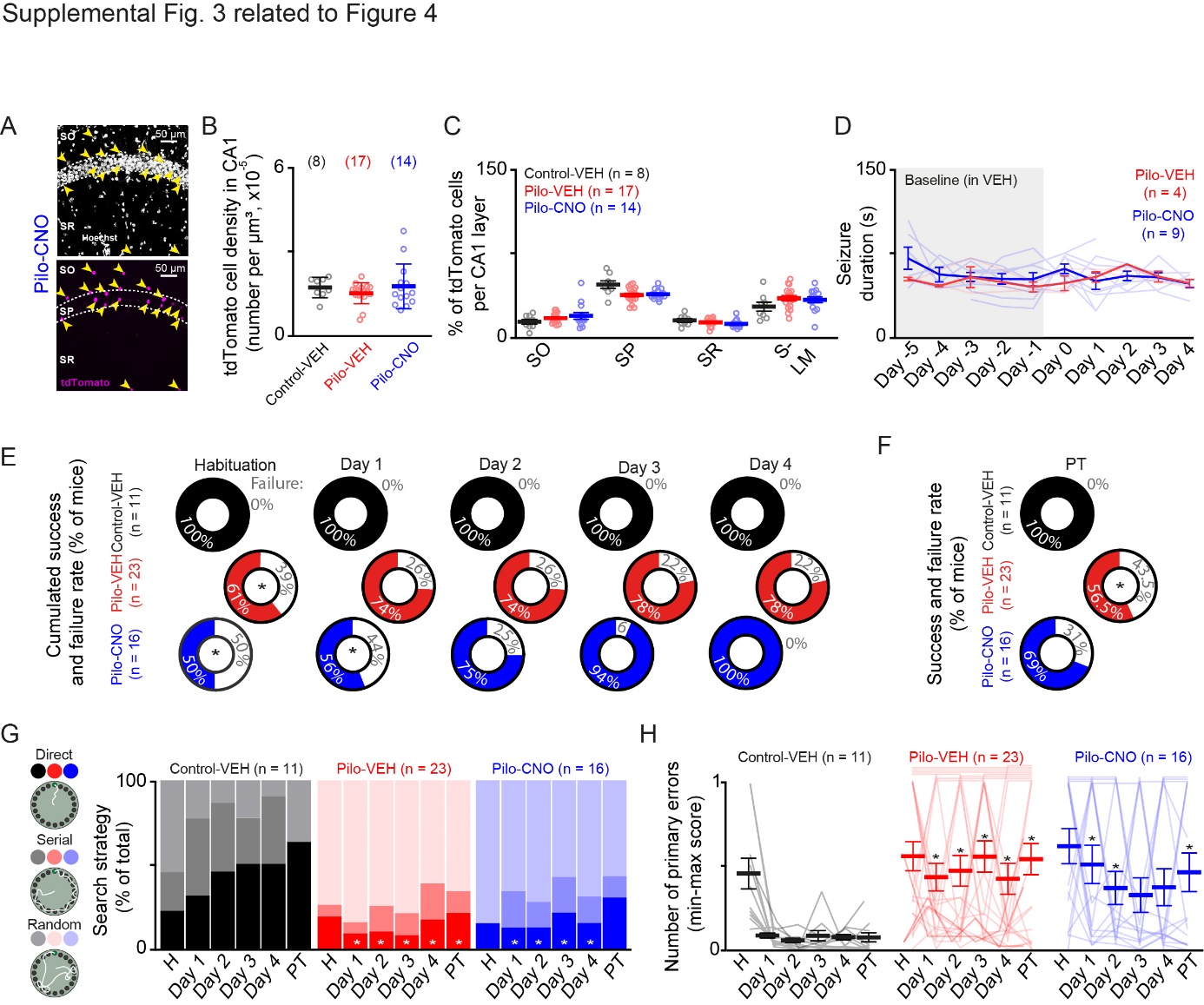

**Supplemental Fig. 3 related to Fig. 4. A.** Representative Hoechst staining (top), as well as tdTomato labelling (bottom) in the CA1 region from a Pilo-CNO mouse infected with AAV9-hDlx-GqDREADD-tdTomato-Fishell-4 in the hippocampus. Neurons with Hoechst and tdTomato co-labelling are indicated by a yellow arrow. **B.** Number of tdTomato-labelled interneurons per µm^3^ of CA1 (Two-tailed Mann-Whitney test: Control-VEH versus Pilo-VEH *P =* 0.2107 and Pilo-VEH versus Pilo-CNO *P =* 0.9454; Two-tailed unpaired t-test: Control-VEH versus Pilo-CNO *P =* 0.8864). **C.** Percentage of tdTomato-labelled interneurons per CA1 layer (Two-way ANOVA experimental condition effect F (2, 36) = 0.8553, *P =* 0.4336). SO: Stratum Oriens, SP: Stratum Pyramidale, SR: Stratum Radiatum, SLM: Stratum Lacunosum-Moleculare. Data are presented as mean ± s.e.m. Each circle represents a given mouse (The number of mice is specified in parentheses). **D.** Duration of ictal events in Pilo-VEH versus Pilo-CNO conditions across days of recording (Two-way ANOVA experimental group effect F (1, 11) = 0.8861 *P =* 0.3667). **E.** Pie charts showing the cumulated percentage of mice that found (success) or not (failure) the location of the escape room at least once during the different days of the BCM acquisition phase. Data are plotted for mice from the following experimental conditions: control mice treated with vehicle (control-VEH, black) and epileptic mice treated with vehicle (pilo-VEH, red) or CNO (pilo-VEH, blue) to specifically elevate or not the activity of hM3D-expressing hippocampal interneurons. The number of mice succeeding or failing to find the escape room at least once in each step and experimental condition was compared using Fisher’s exact test (control-VEH versus pilo-VEH habituation *P =* 0.0172, acquisition days 1-4 *P =* 0.1454, 0.1454, 0.1499, 0.1499; control-VEH versus pilo-CNO habituation *P =* 0.0082, acquisition days 1-4 *P =* 0.0216, 0.1225, 0.9999, 0.9999; Pilo-VEH versus pilo-CNO habituation *P =* 0.5313, acquisition days 1-4 *P =* 0.3115, 0.9999, 0.3703, 0.0660). **F.** Pie charts showing the percentage of mice that found (success) or not (failure) the former location of the escape room during the probe trial. The number of mice succeeding or failing to find the escape room at least once in each step and experimental condition was compared using Fisher’s exact test (control-VEH versus pilo-VEH *P =* 0.0135; control-VEH versus pilo-CNO PT *P =* 0.0598; Pilo-VEH versus pilo-CNO *P =* 0.5166). **G.** Histograms showing the percentage of direct, serial or random exploratory strategies displayed by mice during BCM. The number of mice using a direct versus a serial/random strategy was compared between experimental conditions using Fisher’s exact test (control-VEH versus pilo-VEH habituation *P =* 0.7517, acquisition days 1-4 *P =* 0.0304 0.0034 0.0003 and 0.0087, PT *P =* 0.0256; control-VEH versus pilo-CNO habituation *P =* 0.7131, acquisition days 1-4 *P =* 0.0996 0.0108 0.0423 0.0137, PT *P =* 0.1302 ; Pilo-VEH versus pilo-CNO habituation *P =* 0.7563, acquisition days 1-4 *P =* 0.7100 0.9999 0.7840 0.9999, PT *P =* 0.7110).

**H.** Min-max scores of the number of incorrect holes checked prior to identifying the escape room or the completion of the test (primary errors), in each experimental condition. Data were compared in Control-VEH versus Pilo-VEH groups (Two-way ANOVA experimental group effect F (1, 32) = 16.72 *P =* 0.0003, followed by Sidak’s post hoc test, *P =* 0.9618 0.0022, 0.0010, 0.0003, 0.0064 and 0.0003 for habituation, acquisition days 1-4 and PT, respectively), Control-VEH versus Pilo-CNO groups (Two-way ANOVA experimental group and treatment effect F (1, 25) = 20.20, *P =* 0.0001, followed by Sidak’s post hoc test, *P =* 0.8253, 0.0123, 0.0390, 0.2018 0.0961 and 0.0268 for days 1-4 and PT), and Pilo-VEH versus Pilo-CNO groups (Two-way ANOVA treatment effect F (1, 37) = 1.307, *P =* 0.2603). Straight horizontal lines across the top of the plots indicate that a mouse has scored 1 for two or more consecutive days. Data are presented as mean ± s.e.m unless specified otherwise. Each circle represents a given mouse (The number of mice is specified in parentheses).

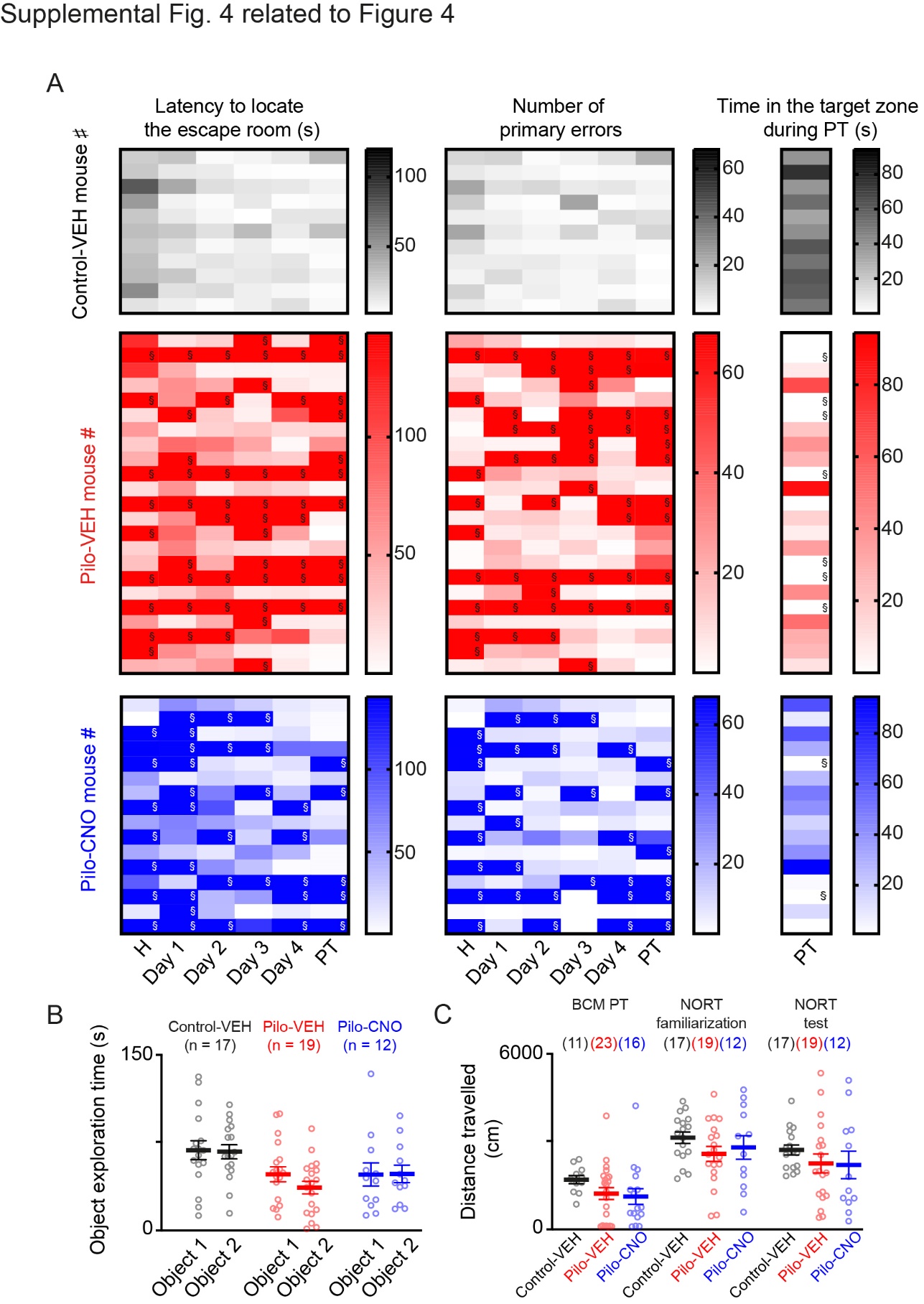

**Supplemental Fig. 4 related to Fig. 4. A.** Latency to locate the escape room (left), number of primary errors (middle) and time in the target zone (right) during the BCM acquisition days (Day 1-4) and/or probe trial (PT) for each Control-VEH (top, black) Pilo-VEH (center, red) or Pilo-CNO (bottom, blue) animals, illustrated with a color scale that represents the minimum to maximum values (maximum: 114 seconds for the latency, 94.43 seconds for the time in the target zone, 68 for the errors). The § symbols indicate cases where no numerical value has been assigned (e.g., the mouse has not found the escape room, has made a low number of errors combined with very little exploration of the arena, ran on the periphery of the table without clear exploration of holes or jumped off the arena, see **Materials and Methods**), a behavior reflecting the worst possible performance, to which we assigned a score of 1 in the min-max normalization. **B.** Time spent exploring each identical object during the NORT familiarization phase, measured in mice from each experimental condition (Two-way ANOVA interaction experimental condition x type of object F (2, 45) = 1.029, *P =* 0.3657). **C.** Distance covered by mice from all three experimental groups during the BCM probe trial as well as NORT familiarization and test phases. Data were compared in Control-VEH versus Pilo-VEH groups (Two-way ANOVA experimental group effect F (1, 54) = 3.900, *P =* 0.0534), Control-VEH versus Pilo-CNO groups (Two-way ANOVA experimental group and treatment effect F (1, 42) = 2.049, *P =* 0.1597), and Pilo-VEH versus Pilo-CNO groups (Two-way ANOVA treatment effect F (1, 52) = 0.0001, *P =* 0.9920). Data are presented as mean ± s.e.m unless specified otherwise. Each circle represents a given mouse (The number of mice is specified in parentheses).

3. **Supplementary references:**

Antunes, M., Biala, G., 2012. The novel object recognition memory: neurobiology, test procedure, and its modifications. Cogn Process 13, 93–110. https://doi.org/10.1007/s10339-011-0430-z

Boileau, C., Deforges, S., Peret, A., Scavarda, D., Bartolomei, F., Giles, A., Partouche, N., Gautron, J., Viotti, J., Janowitz, H., Penchet, G., Marchal, C., Lagarde, S., Trebuchon, A., Villeneuve, N., Rumi, J., Marissal, T., Khazipov, R., Khalilov, I., Martineau, F., Maréchal, M., Lepine, A., Milh, M., Figarella-Branger, D., Dougy, E., Tong, S., Appay, R., Baudouin, S., Mercer, A., Smith, J.B., Danos, O., Porter, R., Mulle, C., Crépel, V., 2023. GluK2 Is a Target for Gene Therapy in Drug-Resistant Temporal Lobe Epilepsy. Annals of Neurology 94, 745–761. https://doi.org/10.1002/ana.26723

Cǎlin, A., Stancu, M., Zagrean, A.-M., Jefferys, J.G.R., Ilie, A.S., Akerman, C.J., 2018. Chemogenetic Recruitment of Specific Interneurons Suppresses Seizure Activity. Frontiers in Cellular Neuroscience 12. https://doi.org/10.3389/fncel.2018.00293

Cohen, S.J., Munchow, A.H., Rios, L.M., Zhang, G., Ásgeirsdóttir, H.N., Stackman, R.W., 2013. The Rodent Hippocampus Is Essential for Nonspatial Object Memory. Current Biology 23, 1685–1690. https://doi.org/10.1016/j.cub.2013.07.002

Dimidschstein, J., Chen, Q., Tremblay, R., Rogers, S.L., Saldi, G.-A., Guo, L., Xu, Q., Liu, R., Lu, C., Chu, J., Grimley, J.S., Krostag, A.-R., Kaykas, A., Avery, M.C., Rashid, M.S., Baek, M., Jacob, A.L., Smith, G.B., Wilson, D.E., Kosche, G., Kruglikov, I., Rusielewicz, T., Kotak, V.C., Mowery, T.M., Anderson, S.A., Callaway, E.M., Dasen, J.S., Fitzpatrick, D., Fossati, V., Long, M.A., Noggle, S., Reynolds, J.H., Sanes, D.H., Rudy, B., Feng, G., Fishell, G., 2016. A viral strategy for targeting and manipulating interneurons across vertebrate species. Nature Neuroscience 19, 1743–1749. https://doi.org/10.1038/nn.4430

Fisahn, A., Pike, F.G., Buhl, E.H., Paulsen, O., 1998. Cholinergic induction of network oscillations at 40 Hz in the hippocampus in vitro. Nature 394, 186–189. https://doi.org/10.1038/28179

Gawel, K., Gibula, E., Marszalek-Grabska, M., Filarowska, J., Kotlinska, J.H., 2019. Assessment of spatial learning and memory in the Barnes maze task in rodents—methodological consideration. Naunyn-Schmiedeberg’s Archives of Pharmacology 392, 1. https://doi.org/10.1007/s00210-018-1589-y

Goirand-Lopez, L., Moulinier, M., Vigier, A., Boileau, C., Carleton, A., Muldoon, S.F., Marissal, T., Crépel, V., 2023. Kainate receptors modulate the microstructure of synchrony during dentate gyrus epileptiform activity. Neurobiology of Disease 185, 106260. https://doi.org/10.1016/j.nbd.2023.106260

Li, P., Geng, X., Jiang, H., Caccavano, A., Vicini, S., Wu, J., 2019. Measuring Sharp Waves and Oscillatory Population Activity With the Genetically Encoded Calcium Indicator GCaMP6f. Frontiers in Cellular Neuroscience 13.

Lim, J.-A., Moon, J., Kim, T.-J., Jun, J.-S., Park, B., Byun, J.-I., Sunwoo, J.-S., Park, K.-I., Lee, S.-T., Jung, K.-H., Jung, K.-Y., Kim, M., Jeon, D., Chu, K., Lee, S.K., 2018. Clustering of spontaneous recurrent seizures separated by long seizure-free periods: An extended video-EEG monitoring study of a pilocarpine mouse model. PLOS ONE 13, e0194552. https://doi.org/10.1371/journal.pone.0194552

Marissal, T., Salazar, R.F., Bertollini, C., Mutel, S., De Roo, M., Rodriguez, I., Müller, D., Carleton, A., 2018. Restoring wild-type-like CA1 network dynamics and behavior during adulthood in a mouse model of schizophrenia. Nature Neuroscience 21, 1412–1420. https://doi.org/10.1038/s41593-018-0225-y

Milosavljevic, N., Cehajic-Kapetanovic, J., Procyk, C.A., Lucas, R.J., 2016. Chemogenetic Activation of Melanopsin Retinal Ganglion Cells Induces Signatures of Arousal and/or Anxiety in Mice. Current Biology 26, 2358–2363. https://doi.org/10.1016/j.cub.2016.06.057

Padmasola, G.P., Friscourt, F., Rigoni, I., Vulliémoz, S., Schaller, K., Michel, C.M., Sheybani, L., Quairiaux, C., 2024. Involvement of the contralateral hippocampus in ictal-like but not interictal epileptic activities in the kainate mouse model of temporal lobe epilepsy. Epilepsia. https://doi.org/10.1111/epi.17970

Peret, A., Christie, L.A., Ouedraogo, D.W., Gorlewicz, A., Epsztein, J., Mulle, C., Crépel, V., 2014. Contribution of Aberrant GluK2-Containing Kainate Receptors to Chronic Seizures in Temporal Lobe Epilepsy. Cell Reports 8, 347–354. https://doi.org/10.1016/j.celrep.2014.06.032

Vigier, A., Partouche, N., Michel, F.J., Crépel, V., Marissal, T., 2021. Substantial outcome improvement using a refined pilocarpine mouse model of temporal lobe epilepsy. Neurobiology of Disease 161, 105547. https://doi.org/10.1016/j.nbd.2021.105547

Zhan, J., Komal, R., Keenan, W.T., Hattar, S., Fernandez, D.C., 2019. Non-invasive Strategies for Chronic Manipulation of DREADD-controlled Neuronal Activity. J Vis Exp 10.3791/59439. https://doi.org/10.3791/59439

Zhang, Y., Rózsa, M., Liang, Y., Bushey, D., Wei, Z., Zheng, J., Reep, D., Broussard, G.J., Tsang, A., Tsegaye, G., Narayan, S., Obara, C.J., Lim, J.-X., Patel, R., Zhang, R., Ahrens, M.B., Turner, G.C., Wang, S.S.-H., Korff, W.L., Schreiter, E.R., Svoboda, K., Hasseman, J.P., Kolb, I., Looger, L.L., 2023. Fast and sensitive GCaMP calcium indicators for imaging neural populations. Nature 615, 884–891. https://doi.org/10.1038/s41586-023-05828-9
